# Noninvasive stimulation of the ventromedial prefrontal cortex modulates rationality of human decision-making

**DOI:** 10.1101/2022.03.15.484390

**Authors:** Thomas Kroker, Miroslaw Wyczesany, Maimu Alissa Rehbein, Kati Roesmann, Ida Wessing, Markus Junghöfer

## Abstract

The framing-effect is a bias that affects decision-making depending on whether the available options are presented with positive or negative connotations. Even when the outcome of two choices is equivalent, people have a strong tendency to avoid the negatively framed option because losses are perceived about twice as salient as gains of the same amount (i.e. loss-aversion). The ventromedial prefrontal cortex (vmPFC) is crucial for rational decision-making, and dysfunctions in this region have been linked to cognitive biases, impulsive behavior and gambling addiction. Using a financial decision-making task in combination with magnetoencephalographic neuroimaging, we show that excitatory compared to inhibitory non-invasive transcranial direct current stimulation (tDCS) of the vmPFC reduces framing-effects while improving the assessment of loss-probabilities, ultimately leading to increased overall gains. Behavioral and neural data consistently suggest that this improvement in rational decision-making is predominately a consequence of reduced loss-aversion. These findings recommend further research towards clinical applications of vmPFC-tDCS in addictive disorders.

## Introduction

We humans like to believe that our decision-making behavior follows rational considerations. We think that as rational people we can conscientiously analyze the consequences of available options by weighing their probabilities and comparing the results to finally choose the optimal option that maximizes our gain and minimizes our loss. However, in pioneering studies leading to the development of the ‘prospect theory’, Kahneman and Tversky showed that humans often tend to rely on heuristics and other cognitive shortcuts, instead of employing cognitively demanding evaluations (Kahneman and Tversky, 1979; Tversky and Kahneman, 1992). Remarkable deviations from the behavior of an ideal rational agent (‘homo economicus’) are direct consequences of a strong loss-aversion, as humans rate losses twice as salient as gains of the same amount (Kahneman and Tversky, 1979; Tversky and Kahneman, 1992). This cognitive bias of loss-aversion likely roots in an asymmetric evolutionary pressure on losses and gains: “when survival is uncertain, marginal losses prove more critical for reproductive success than marginal gains” (McDermott et al., 2008). Loss-aversion has been used to explain a wide range of economic behaviors that compromise rationality such as the sunk-cost effect (Arkes and Blumer, 1985; i.e. the belief that previous investments and even losses justify further expenditures) or the status-quo bias (Kahneman, Knetsch and Thaler, 1991; i.e. changes from the status-quo are expected as rather negative). A well-studied example to illustrate further far-reaching consequences of loss-aversion is the so called framing-effect, where presenting either positive or negative connotations of an option can bias decisions: For instance, when people are asked to either opt for a treatment A which would *save* 400 of overall 600 infected patients or for treatment B with which 200 of 600 patients would *die*, a vast majority would prefer treatment A, which is positively framed, but still identical to B in terms of outcomes (Tversky and Kahneman, 1981).

The impact of the framing-effect and reactions to odds — and therefore the modulation of rational decision-making by any kind of intervention — can nicely be investigated in gambling studies that can consider gains and losses simultaneously. For example, if participants have to decide between ‘*lose* 60ct of 100ct’ or ‘gamble for 100ct with 20% probability’ they would typically avoid the negatively (‘*lose’*) framed option and gamble for the whole amount, despite a higher expected safe residual gain (40 cent) in contrast to the expected gain (20 cent) of the gambling decision. Inversely, most participants would choose the positively framed ‘*keep* 40ct of 100ct’ instead of ‘gamble for a 100ct with 80% probability’, despite the expected gain of the gambling option (80 cent) would be higher than the safe gain of 40 cents.

Studies investigating the underlying neural correlates of loss-aversion and the resulting framing-effect in gambling studies consistently identified the ventromedial prefrontal cortex (vmPFC) as a cardinal player. In fact, neural activity in the vmPFC decreased with increasing potential losses and the strength of this association predicted individual loss-aversion (Tom et al., 2007). Consistently, individuals with increasing susceptibility to the framing-effect showed decreased activity in the vmPFC (Martino et al., 2006). The finding that patients with vmPFC lesions feature an increased framing-effect relative to patients with other lesions and healthy controls (Pujara et al., 2015) adds causality to this correlational evidence.

Another aspect of rational decision-making that is dependent on the vmPFC is the inhibition of impulsive and short-sighted choices. Patients with vmPFC lesions for instance show a distinct insensitivity to consequences of decision-making and are primarily guided by immediate prospects (Kahneman and Tversky, 1983). These patients also show increased risk-taking and bet more money in gambling studies than healthy controls (Studer et al., 2015). This causal link between vmPFC dysfunctionality and irrational economic behavior is further supported by the finding that the reduction of rational economic decision-making with aging correlates with gray matter volume reduction in ventral PFC regions (Chung et al., 2017).

Since reduced vmPFC activity and vmPFC dysfunctions have been associated with irrational decision-making based on loss-aversion, as indexed by an increased susceptibility to framing-effects and poorer assessments of the odds, it appears tempting to assume that excitatory vmPFC stimulation (i.e. increasing vmPFC excitability) might have mitigating effects on these biases and might improve rational decision-making. Because the vmPFC is an almost unique structure in coding gains with activity increase and losses with activity decrease (Lindquist et al., 2016; Tom et al., 2007), the vmPFC recommends itself as ideal target for valence specific interventions via brain stimulation (i.e. modulation of valence biases such as loss-aversion). In fact, a series of fMRI and MEG studies from our lab recently revealed that excitatory stimulation of the vmPFC attenuated behavioral and neural negativity-biases to emotional scenes and emotional facial expressions in healthy participants (Junghofer et al., 2017; Winker et al., 2020, 2019, 2018).

Here we combined the financial gambling paradigm developed by Kahneman and Tversky (1983) - and adopted by De Martino and coworkers (2006) as well as Pujara and colleagues (2015) - with the vmPFC stimulation and MEG neuroimaging approach as used in our previous studies (Junghofer et al., 2017; Winker et al., 2020, 2019, 2018). Within this paradigm, participants decided to either accept a safe amount of an initial stake or gamble for the whole amount with variable risks to win or lose. The safe option was either framed in a positive or negative way with identical net gains for both frames. The probability of taken risks thus informed about the susceptibility to the framing-effect and the consideration of odds which here were operationalized as indices of rational decision-making. Participants finally received feedback about gains and losses. The participants’ evaluation of gains and losses informed about the strength of the framing-effect in this feedback phase which was used as further index of rational decision-making.

Directly preceding the financial gambling task, one half of the participants received excitatory vmPFC stimulation at a first session and inhibitory vmPFC stimulation at a second session while the other half received the reversed order. We hypothesized that vmPFC-excitation by transcranial Direct Current Stimulation (tDCS) would result in a decreased loss-aversion. In contrast to inhibitory stimulation, this should be reflected in an attenuated framing-effect (i.e. a decreased tendency to gamble in the ‘loss-frame’ and thus more rational decision-making) as well as an improved consideration of odds and ultimately higher gains.

Investigations of event related potentials (ERPs) of gain and loss processing during gambling tasks have consistently identified ERP components at mid-latency to late time intervals (around 250ms after feedback) and at central to frontal-central scalp regions (Gehring and Willoughby, 2002; Hajcak et al., 2006; Yeung et al., 2005). While these components typically reflect responses to both positive and negative feedback, stronger reactions to losses as compared to gains, most presumably reflecting loss-aversion, have been identified in various ERP studies (Hajcak et al., 2006; Yeung et al., 2005). Thus, MEG correlates of reduced loss-aversion and attenuated framing after excitatory stimulation were expected to occur at prefrontal regions and mid-latency to late time intervals. Since previous research has shown that the vmPFC mediates rationality by inhibiting maladaptive responses (i.e. irrational loss-aversion or choosing high-risk options; Boes *et al*., 2009; Manuel, Murray and Piguet, 2019) we expected that the neural data would reflect an enhanced ability of prefrontal brain regions to inhibit loss-aversion (in the decision-phase and feedback-phase) and thus maladaptive choices (in the decision-phase) following excitatory compared to inhibitory stimulation.

With the focus on effects of vmPFC stimulation on rational decision-making and feedback-processing, behavioral and neural effects which were modulated by the stimulation are reported in the following results section. For main effects and further interactions please consult the supplementary material (SM).

## Results

### Non-invasive vmPFC-tDCS modulates rational decision-making

We applied a within-subjects design so that each participant received excitatory and inhibitory stimulation of the vmPFC on two different days with at least 48 hours in between (Methods Fig. 1). The tDCS was optimized for targeting the vmPFC and minimizing the impact on other brain regions (Methods Fig. 2). In both excitatory and inhibitory conditions participants were stimulated with a current strength of 1.5mA over 10 minutes and were unable to differentiate between stimulation conditions (see SM2.1). Immediately after stimulation, participants performed the gambling task, where gambling behavior and event-related magnetic fields were measured. At the beginning of each trial, participants received an initial amount (‘gambling stake’), which was varied (25ct, 50ct, 75ct, 100ct) to enhance the credibility of the gambling paradigm (see Fig. SM2). Participants were asked to either keep a smaller amount or to gamble for the full amount. The ‘keep’ option was either framed as a gain (Fig. 1A; gain-frame: receive a smaller but safe amount) or framed as a loss (Fig. 1A; loss-frame: subtraction of a smaller safe amount). Although the final monetary amounts were equivalent in both frames, participants chose the ‘gamble’ option in the loss-frame and accepted the safe smaller amount in the gain-frame more often, replicating the framing-effect(Kahneman and Tversky, 1983) (Fig. 1B; *z* = 2.64, *p* = 0.008). Next, we specifically tested the hypothesis that noninvasive vmPFC stimulation modulates the framing-effect as index of rational decision-making. Indeed, a logistic regression with the predictors of stimulation (excitatory, inhibitory) and frame (gain-frame, loss-frame) revealed a significant interaction (*z* = 2.19, *p* = 0.029) modulating the decision (keep, gamble). Importantly and supporting our hypothesis, excitatory vmPFC stimulation resulted in a significantly reduced framing-effect compared to inhibitory stimulation (Fig. 1B; *χ²* = 14.74, *p* < 0.001). Furthermore, excitatory stimulation significantly reduced the proportion of gambling choices in the loss-frame (*χ²* = 12.24, *p* < 0.001) but not in the gain-frame (*χ²* = 0.086, *p* = 0.769) supporting the hypothesis that reduced loss-aversion after excitatory stimulation underlies the attenuated framing-effect.

**Figure 1.**
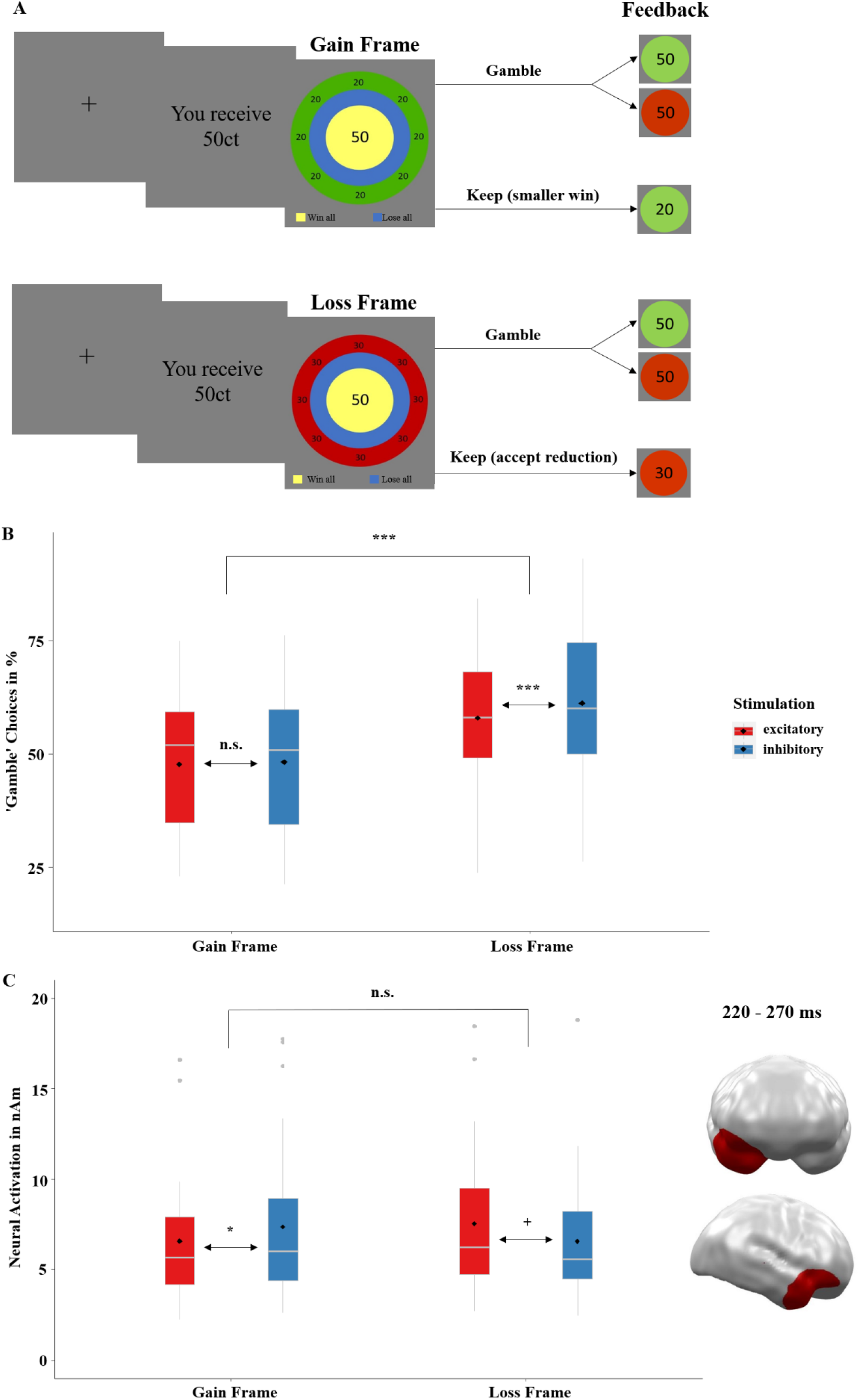
**A**. Course of a single trial in the gambling task adapted from Martino and colleagues (2006). Each trial began with a fixation cross presented for 500ms, followed by the presentation of the ‘game stake’ of 25, 50, 75 or 100 cents. The subsequent ‘choice stimulus’ reminded participants of the initial amount (center), the chance to win or the risk of losing when choosing the ‘gamble’ option (20%, 40%, 60%, 80%, based on the relative sizes of the blue and yellow circles), and the frame when choosing the safe ‘keep’ option (green and red outer ring for gain and loss frames, respectively). After choosing the ‘keep’ or ‘gamble’ option, feedback on the win or loss was given via green (win) and red (loss) circles, with the amount depicted in the center. Stimuli were placed centrally to minimize eye movements and related MEG artifacts. MEG correlates of neural activity evoked by the choice and the feedback stimuli were analyzed. **B**. Proportion of ‘gamble’ choices for gain- and loss-framed trials in percentage. An ideal rational agent would have chosen the gain-frame option and the loss-frame option in equal frequency, as both resulted in identical wins or losses. However, replicating a strong deviation from rationality, participants chose the risky ‘gamble’ option in the loss-framed condition much more often. Importantly, this framing-effect (i.e., a preference for gambling in the loss frame compared to in the gain frame) was stronger after inhibitory than excitatory stimulation of the vmPFC. Convergent with the idea that vmPFC excitation modulates loss-aversion, the effect of exciting the vmPFC was highly significant in the loss frame but not in the gain frame. **C**. Significant spatio-temporal cluster in right anterior temporal/orbitofrontal areas featuring an interaction effect of stimulation by frame. The relatively greater neural activation in response to the loss frame after excitatory tDCS suggests that vmPFC excitation results in more elaborate inhibition of the loss-frame processing than in vmPFC inhibition, such that excitation leads to a reduced saliency of the loss condition and, eventually, a reduced framing-effect, as seen in B. The location of this cluster agrees well with results of the Martino et al. study (2006), in which enhanced activity in the vmPFC as well as in right OFC regions was associated with more rational decision-making. Topographies of effects observed in L2-MNE were projected on standard 3D brain models for visualization. Boxplots indicate means (black dot), medians (grey line) and lower and upper quartiles. Asterisks indicate significance levels: + < 0.1, * < 0.05, ** < 0.01, *** < 0.001.

To uncover underlying neural correlates of the framing-effect and its interaction with stimulation, we computed a 2×2×2 repeated-measures ANOVA on source-localized MEG data (the dependent variable; see SM1.4) with the factors stimulation (excitatory, inhibitory), frame (gain-frame, loss-frame) and decision (keep, gamble). The interaction of stimulation by frame was significant at the junction of right anterior temporal and right orbitofrontal regions in a mid-latency time interval between 220 and 270ms (*p*-cluster = 0.023; Fig. 1C). Post-hoc t-tests of this interaction revealed less activation in the gain-frame after excitatory compared to inhibitory stimulation (*t* = 1.77, *p* = 0.044), while activation in the loss-frame showed the opposite trend-significant effect (*t* = -1.65, *p* = 0.055). The relatively greater neural response to the ‘loss-frame’ after excitatory tDCS suggests an improved inhibition of loss-aversion, most probably leading to a reduced salience of losses and eventually resulting in an attenuated framing-effect as seen in the behavioral stimulation effects. The location of the cluster nicely converges with fMRI findings of Martino and coworkers (2006), who showed that, in addition to vmPFC areas, enhanced activity at right orbitofrontal cortex regions (OFC) was also associated with more rational decision-making as reflected by reduced framing-effects.

Next, we aimed to investigate whether the more rational decision-making following excitatory versus inhibitory vmPFC stimulation as shown above in the ‘keep’ option generalized to decision-making depending on the odds (i.e. when choosing the ‘gamble’ option). Here, the risk-to-lose or the chance-to-win respectively was varied in steps of 20%, 40%, 60% and 80%, indicated by the size relation of the inner blue and yellow circles in both frames (Fig. 2A). Of course, participants increasingly avoided gambling with increasing risk-to-lose (*z* = -9.60, *p* < 0.001). But importantly, the effect of risk-to-lose on gambling behavior was significantly modulated by stimulation (risk-to-lose by stimulation: z = 6.60, p < 0.001; Fig. 2B). Post-hoc tests revealed more risky-choices in the two low-risk conditions (20%: *χ*² = 35.81, *p* < 0.001 and 40%: *χ*² = 4.81, *p* = 0.028) but fewer risky-choices in the two high-risk conditions (60%: *χ*² = 37.32, *p* < 0.001 and 80%: *χ*² = 40.97, *p* < 0.001) after excitatory compared to inhibitory stimulation. A representation of the averaged winnings achieved in the game across all choices (‘keep’ or ‘gamble’; Fig. 2C) elucidates the consequences of the participant’s behavior. A constant choice of the ‘keep’ option would have resulted in averaged winnings of 25 cents across all risk conditions (Fig. 2C dotted line). A constant choice of the ‘gamble’ option would have resulted in averaged winnings of 50 cents at 20% risk, 37.5 cents at 40% risk, 25 cents at 60% risk and 12.5 cents at 80% risk (Fig. 2C dashed lines). Thus, participants behaved quite rational as they almost reached the maximal win for all risk conditions. However and importantly, a t-test across all trials (t(11519) = 2.23; p = 0.026) revealed that after excitatory vs. inhibitory stimulation averaged wins were significantly higher. Post-hoc t-tests showed significantly higher winnings in the lowest 20% (*t* = 1.74, *p* = 0.041) and predominately in the highest 80% (*t* = 2.95, *p* = 0.002) risk conditions after excitatory compared to inhibitory stimulation. Averaged winnings did not differ in the 40% condition (*t* = 0.62, *p* = 0.267) and could not differ in the 60% risk-to-lose conditions because both ‘keep’ and ‘gamble’ choices resulted in an identical amount of 25ct.

**Figure 2.**
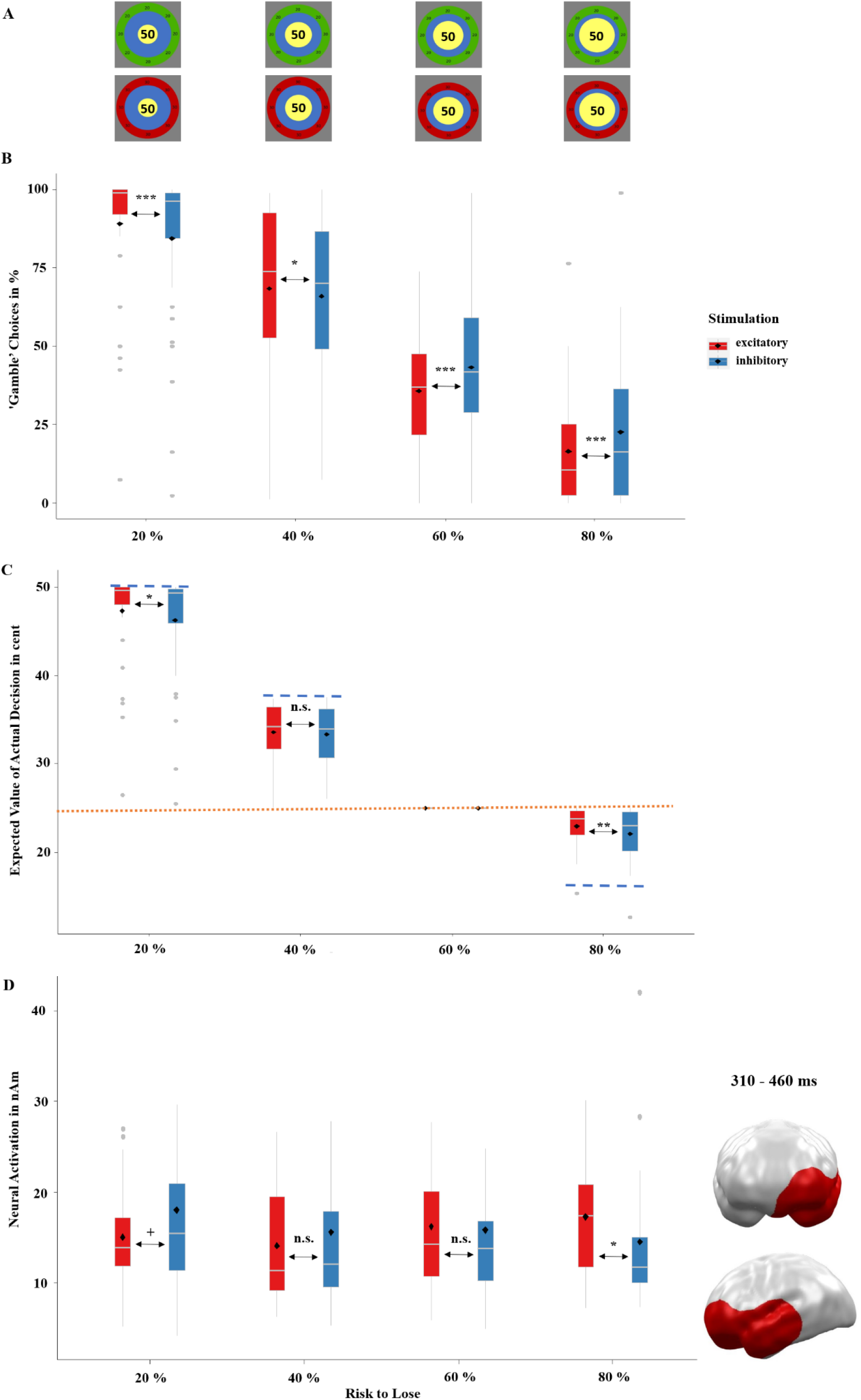
**A**. The relative risk-to-lose percentages or chance-to-win percentages, respectively, when choosing the ‘gamble’ option was based on the relative sizes of the blue and yellow inner circles of the choice stimulus. **B**. Proportion of ‘gamble’ choices in percentage depending on the respective risk-to-lose. After excitatory stimulation, participants gambled more often at the lower risk-to-lose conditions (20% and 40%) but avoided gambling more often at higher risk-to-lose conditions (60% and 80%). **C**. Mean expected values of actual choices in cents depending on the respective risk-to-lose conditions. The mean expected outcome for the ‘keep’ option averaged across all initial amounts was 25 cents (dotted orange line in C). The mean expected outcome of the ‘gamble’ option across all initial amounts was 50 cents at 20% risk, 37.5 cents at 40% risk, 25 cents at 60% risk and 12.5 cents at 80% risk (dashed blue lines in C). Concluding, to maximize the expected value (i.e., their overall winnings) participants should have always chosen the ‘gamble’ option in the 20% and 40% risk conditions and should have always chosen the ‘keep’ option in the 80% risk condition (in the 60% risk condition, the ‘keep’ and ‘gamble’ options led to identical averaged wins). Without stimulation, participants performed already quite rationally, as they almost reached the maximal wins for each risk-to-lose condition. However, in the low-risk 20% condition and the high-risk 80% condition, participants reached significantly higher winnings after excitatory compared to inhibitory stimulation. As the ‘keep’ and ‘gamble’ choices both resulted in 25 cents for the 60% condition, the respective expected values were always identical and are shown for clarity reasons only. **D**. Significant spatio-temporal cluster at left prefrontal and anterior temporal areas featuring an interaction effect of stimulation by risk-to-lose condition. Integration of these neural responses with the behavioral results (Fig. 2B&C) suggests that excitatory (versus inhibitory) stimulation gave participants a greater ability to inhibit inadequate risky behavior in the high-risk 80% condition, while it reduced the inhibition of risky behavior (i.e., risky behavior was facilitated) in the low-risk 20% and semi-low-risk 40% conditions. Thus, convergent with the behavioral effects, this neural pattern suggests that excitatory compared to inhibitory stimulation facilitates rational decision-making toward the expected value (Fig. 2C). Topographies of effects observed in L2-MNE were projected on standard 3D brain models for visualization. Boxplots indicate means (black dots), medians (grey lines) and lower and upper quartiles. Asterisks indicate significance levels: + < 0.1, * < 0.05, ** < 0.01, *** < 0.001.

On the neural level, we performed a 2×4 repeated-measures ANOVA with the factors stimulation (excitatory, inhibitory) and risk-to-lose (20%, 40%, 60%, 80%) on source-localized MEG data (dependent variable). This analysis revealed an interaction of stimulation by risk-to-lose between 310 and 460ms (Fig. 2D) at left anterior temporal and left orbitofrontal regions (*p*-cluster = 0.025) and thus later but laterally symmetric to the above reported interaction of stimulation by frame (Fig. 1C). Post-hoc tests of neural activity within this cluster revealed that this interaction was driven by the lowest and highest risk conditions (20%: *t* = -1.56, *p* = 0.065; 80%: *t* = 1.85, *p* = 0.038), while the medium risks, did not show significant effects of stimulation (40%: *t* = -0.82, *p* = 0.209; 60%: *t* = 0.23, *p* = 0.409). Convergent with the behavioral effects, this neural pattern suggests that excitatory compared to inhibitory stimulation facilitates rational decision-making towards maximized winnings.

Taken together, the behavioral data consistently indicates a modulation of decision-making towards increased rationality after excitatory compared to inhibitory vmPFC-stimulation. The reduced framing-effect and greater ability to estimate risks is mirrored by the neural data providing relatively enhanced inhibition of loss-aversion and high-risk options after excitatory versus inhibitory stimulation.

### Non-invasive vmPFC-tDCS modulates loss-aversion in feedback-processing

After the participant’s choice, the feedback on win or loss was indicated by green and red circles with the amounts of the wins or losses in the middle, respectively (Fig. 3A). Finally, participants rated their subjective hedonic valence and emotional arousal in response to each outcome. Having established that neurostimulation significantly modulated the rationality of decision-making, here we aimed to determine the effects of tDCS on rational feedback-processing in particular on its modulation of loss-aversion. We addressed this question by computing a mixed effects linear regression with the predictors stimulation (excitatory, inhibitory), outcome (gain, loss) and decision (keep, gamble). A main effect of outcome (*t* = -25.08, *p* < 0.001; Fig. 3A) reflected the trivia that gains were rated more positive than losses. Importantly, feedback evaluations were overall (across keep and gamble decisions; Fig. 3A&B) rated more positive after excitatory than inhibitory stimulation (*t* = 2.99, *p* = 0.003). A main effect of decision (*t* = -5.16, *p* < 0.001; keep > gamble) was mainly driven by a less negative evaluation of losses in the ‘keep’ condition after excitatory stimulation which will be further discussed below. While stimulation did not affect the factors decision and outcome alone (stimulation by decision: *t* = -0.17, *p* = 0.867; stimulation by outcome: *t* = 1.65, *p* = 0.245), the three-way interaction was significant (stimulation by decision by outcome: *t* = 5.33, *p* < 0.001). Post-hoc repeated-measures ANOVAs that were conducted separately for the ‘keep’ and ‘gamble’ conditions revealed a significant interaction of stimulation by outcome in the ‘keep’ condition (*F*(1, 35) = 11.48, *p* = 0.001; Fig. 3A), while the respective interaction was insignificant in the ‘gamble’ condition (*F*(1, 35) = 1.79, *p* = 0.190; Fig. 3B) indicating that the three-way interaction was mainly driven by the ‘keep’ condition. To further elucidate the influence of outcome in the three-way interaction and the effect of stimulation on rational decision-making, we calculated the difference of gain-ratings minus loss-ratings after ‘keep’ and ‘gamble’ decisions and calculated t-tests comparing excitatory and inhibitory stimulation. With respect to modulations of the framing-effect, the difference of gain-ratings minus loss-ratings in the ‘keep’ condition (Fig. 3A) was of major interest since both options held the same monetary value while monetary outcomes were very different in the ‘gamble’ condition (Fig. 3B). As predicted, the ‘framing-difference’ was smaller following excitatory compared to inhibitory tDCS (*t*(35) = -3.14, *p* = 0.003), reflecting less loss-averse feedback-processing, while the corresponding t-test in the ‘gamble’ condition was insignificant (*t*(35) = 1.64, *p* = 0.111).

**Figure 3.**
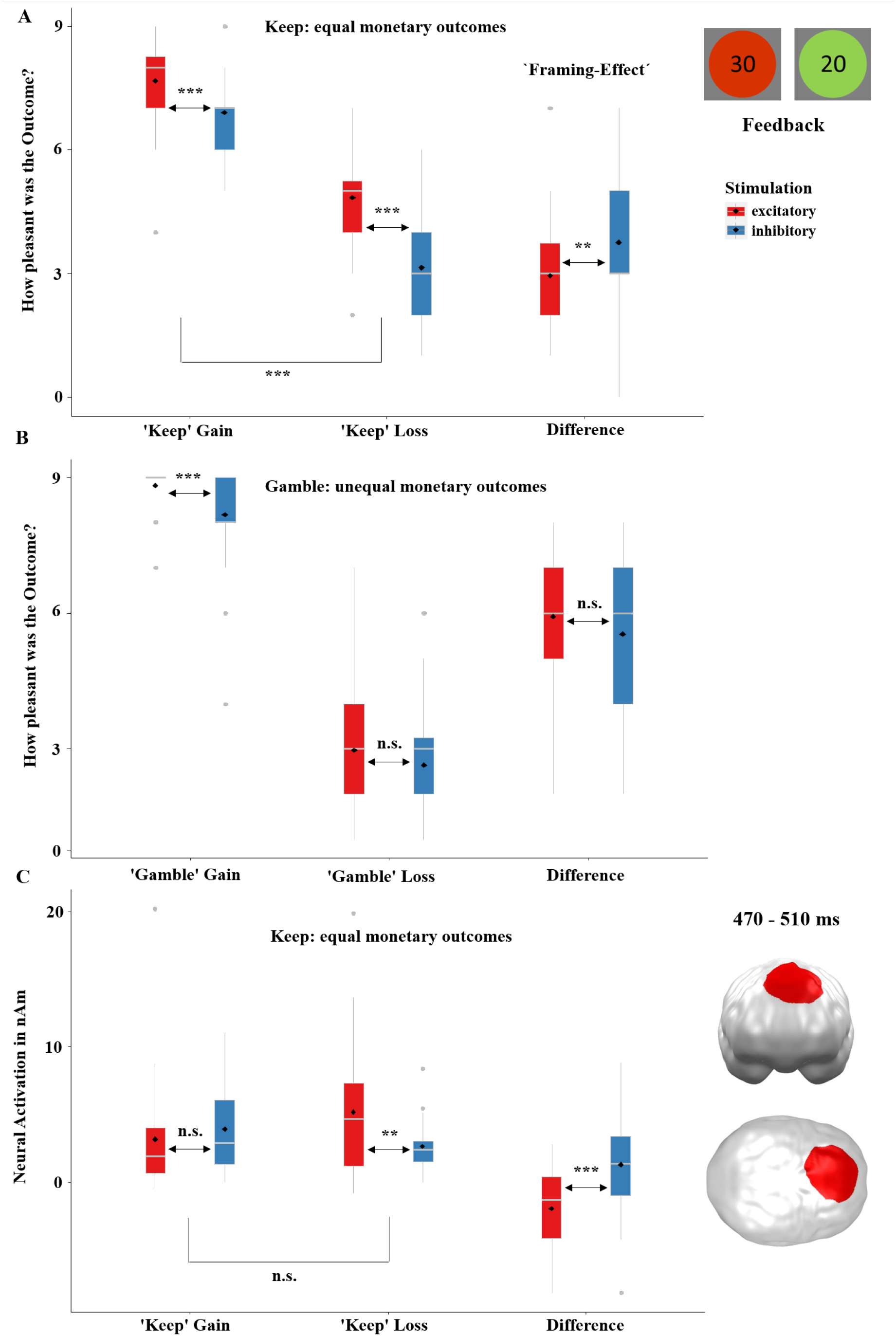
**A**. Rated hedonic valence (pleasantness) on a SAM scale - 1 (most negative) to 9 (most positive) - in the ‘keep’ condition with equal monetary outcomes. **B**. Rated hedonic valence in the ‘gamble’ condition with unequal monetary outcomes. Excitatory stimulation led to an overall (across keep and gamble option) more positive feedback evaluation and resulted in a relatively reduced framing-effect (the difference between gain and loss ratings in the ‘keep’ option, A). Thus, excitatory stimulation led to an attenuated loss-aversion-bias and to more rational (i.e., less loss averse) feedback-processing. **C**. Significant spatio-temporal cluster in the dorsomedial prefrontal cortex featuring a significant effect of stimulation revealed by a t-test employing the difference of gain minus loss in the relevant framing ‘keep’ option. The greater activation in response to ‘keep’ losses following excitatory compared to inhibitory stimulation suggests that excitation stimulation helped inhibit negative feedback, leading to a reduced negative evaluation of losses in the keep option (A) and eventually to a more rational (i.e., less loss averse) evaluation of feedback stimuli. Topographies of effects observed in L2-MNE were projected on standard 3D brain models for visualization. Boxplots indicate means (black dots), medians (grey lines) and lower and upper quartiles. Asterisks indicate significance levels: + < 0.1, * < 0.05, ** < 0.01, *** < 0.001.

To investigate the neural responses to the feedback stimuli, we performed a 2×2×2 ANOVA employing the factors stimulation (excitatory, inhibitory), decision (keep, gamble) and outcome (gain, loss). In a repeated-measures ANOVA, this did not reveal a significant main effect of stimulation nor significant interactions with the factor stimulation. This was most likely due to the extremely strong main effect of outcome (loss >> gain) - reflecting the strong loss-aversion since losses are usually rated twice as salient as gains - that explained a large part of variance (see SM2.4.3). Nevertheless, we tested our specific hypothesis regarding the modulation of framing-effects via tDCS by subtracting gains from losses in the ‘keep’ option (i.e. the relevant comparison to evaluate the effect of stimulation on framing) and used the resulting difference for a t-test (excitatory versus. inhibitory). This revealed a significant cluster (*p-*cluster = 0.045) in the dorsomedial prefrontal cortex in a late time interval between 470 and 510ms (Fig. 3C). Post-hoc tests of neural activity within this cluster revealed greater activations in response to losses after excitatory compared to inhibitory stimulation (*t* = 2.79, *p* = 0.010) while stimulation did not modulate gain trials (*t* = -0.79, *p* = 0.437). The increased responses to losses in the ‘keep’ condition following excitatory stimulation suggest facilitated inhibition of loss-aversion leading to less negative evaluations of losses (Fig. 3A) and eventually more rational evaluation of feedback stimuli.

In summary, convergent to effects in the decision-making phase, the analysis of feedback-processing also revealed that excitatory compared to inhibitory stimulation reduced behavioral loss-aversion and enhanced inhibition of loss-aversion in neural measures.

## Discussion

We investigated the contribution of the vmPFC on rational decision-making and feedback-processing by increasing or reducing its excitability via non-invasive tDCS. We found evidence for a causal role of vmPFC-activity in rational decision-making on both behavioral and neural levels. Excitatory stimulation induced a more adaptive decision-making, which was reflected by reduced loss-aversion that resulted in less susceptibility to the framing-effect, a greater likelihood to choose the option providing the higher expected value and a decreased tendency to risk larger amounts (SM2.3.2). The neural data supported this pattern, suggesting an increased ability of bilateral ventral prefrontal cortex regions to inhibit maladaptive choices (reduction of loss-aversion and less frequent choice of options with a high risk-to-lose) and thus enable more adaptive and less impulsive gambling behavior after excitatory stimulation. The same applied to the feedback-processing, as reduced loss-aversion with less susceptibility to the framing-effect was present after excitatory stimulation in both, behavioral and neural data. Furthermore, excitatory stimulation resulted in an overall reduction of the loss-aversion-bias for feedback-processing. Overall, our results show that non-invasive brain stimulation of the vmPFC can influence the rationality of human decision-making and feedback-processing as seen on both behavioral and neural levels.

Remarkably, we could show that vmPFC-inhibition led to increased susceptibility to the framing-effect. As seminal studies have previously shown (Kahneman and Tversky, 1983, 1979), participants exhibited more risk-taking behavior in the ‘loss-framed’ option, compared to the ‘gain-framed’ option. This bias was further increased by inhibitory compared to excitatory stimulation, as participants featured even more risk-taking behavior in the ‘loss-framed’ option after vmPFC-deactivation (Fig. 1B). The importance of vmPFC-activation to integrate frames into decisions has been found before (Deppe et al., 2005) and is present in our neural data as well (Fig. 1C). We found greater activity in response to the ‘loss-frame’ following excitatory stimulation in the right OFC replicating a correlation of right OFC-activity and rational decision-making in the fMRI study from Martino and coworkers (2006). Greater OFC-activation to the ‘loss-frame’ after excitatory stimulation suggests an improved inhibition of a potentially irrational loss-aversion. In contrast, inhibitory stimulation could even strengthen this non-adaptive tendency (i.e. intensifying loss-aversion) and promote a more impulsive decision for the ‘gamble’ option, although the expected value may be higher in the ‘loss-framed’ option. Accordingly, a study employing delayed reward found that excitatory tDCS of the vmPFC improved the ability to wait for rewards and reduce impulsivity accordingly (Manuel et al., 2019).

Second, vmPFC-excitation decreased risk-taking behavior, when the risk of losing was high and increased risk-taking, when the chance to win was high, overall enhancing the expected value of decisions and the overall wins compared to inhibitory stimulation (Fig. 2B&C). This is consistent with previous findings showing that patients with vmPFC-lesions are less able to assess the probability of gains/losses and consequently win less money than healthy controls (Studer et al., 2015). The spatiotemporal MEG cluster of stimulation by risk-to-lose overlapping the vmPFC (Fig. 2D) may illuminate the underlying neural mechanism: after inhibitory stimulation an increasing risk-to-lose came along with decreasing prefrontal activity, suggesting that vmPFC-inhibition reduced the ability to suppress maladaptive choices. By contrast, excitatory tDCS facilitated this capability in participants as indicated by increasing prefrontal activity with increasing risk to lose, in turn leading to more adaptive decision-making. This result can be underlined with findings of an fMRI study, where greater activity in medial prefrontal areas occurred when risk anticipation was necessary (Fukui et al., 2005). Concluding, our results combined with previous findings strongly suggest that ventro-medial prefrontal activity is responsible for risk assessment and that this process can be modulated by non-invasive tDCS.

Another important finding is the increased tendency to gamble with increasing ‘game stakes’ after inhibitory stimulation, while risk-taking behavior did not change after vmPFC-excitation (see SM2.3.2). This is consistent with the finding that vmPFC-lesioned patients typically bet more money than healthy participants (Studer et al., 2015). Additionally, pathological gamblers show vmPFC-hypoactivation and its strength was correlated with gambling severity (Reuter et al., 2005). Combined with the finding that excitatory stimulation induced less risk-taking behavior overall and specifically in high-risk situations, these findings suggest that excitatory vmPFC-tDCS might be a promising add-on treatment option for patients suffering from gambling addiction. It should be noted, that the reported behavioral and neural effects occurred after only one rather short and mild excitatory/inhibitory tDCS (far below the thresholds considered to be safe; Sparing and Mottaghy, 2008). Thus, follow-up clinical tests on add-on therapy effects have the option to use more sessions, longer durations and greater intensities of excitatory stimulation.

Now considering feedback-processing we can further elaborate the idea that the vmPFC modulates loss-aversion and, thus, is important for more rational (i.e. less biased) feedback-processing and can be modulated through tDCS. We identified that the significant interaction of stimulation by decision by outcome (Fig. 3A and 3B) was driven by a greater framing-effect after inhibitory stimulation. Thus, as in decision-making, more rational feedback-processing (i.e. decreased susceptibility to framing) occurred after excitatory compared to inhibitory stimulation. This can also be explained by a model that sees the vmPFC as a region tracking the (financial) value of a decision providing the basis for learning from reward and punishment. This function is implied by lesion (Liu et al., 2011) and functional imaging (Pujara et al., 2015; Tom et al., 2007) studies, which suggest that the vmPFC is also responsible for evaluating decisions and guiding future behavior by these experiences. The relevance of the vmPFC for learning and prediction is underlined by a study on cocaine addicts (Parvaz et al., 2015), in which the so called feedback-negativity, an ERP-component that is predominately modulated by vmPFC-activity (Carlson et al., 2011), was examined. Cocaine addicts exhibited an impaired feedback-processing in response to losses (Parvaz et al., 2015), indicating their inability to learn from losses possibly related to vmPFC-hypoactivation eventually resulting in compromised predictions.

More positive feedback ratings overall following excitatory vmPFC-tDCS also dovetail with our previous findings of a relative reduced negativity-bias of emotional face and emotional scene processing after excitatory compared to inhibitory stimulation (Junghofer et al., 2017; Winker et al., 2020, 2019, 2018). Indeed, losses in the ‘keep’ option in particular were rated as less negative after excitatory compared to inhibitory stimulation, which fits with the hypothesis that vmPFC-excitation reduces loss-aversion. Convergent, losses in the ‘keep’ option also evoked stronger neural responses after excitatory compared to inhibitory stimulation (Fig. 3C). Like the strengthened neural responses to loss-frames in right anterior temporal/orbitofrontal areas (Fig. 1C) after excitatory stimulation - which was interpreted as improved inhibition of a loss-aversion-bias - the neural responses to ‘keep’ losses in dorsomedial prefrontal areas also suggest an enhanced inhibition of loss processing eventually leading to a less negative evaluation of ‘keep’ loss outcomes as seen in figure 3A. In fact, dorsolateral and dorsomedial prefrontal areas are typically active during implicit (Roesmann et al., 2020) and explicit (Ochsner and Gross, 2005) suppression of negative stimulus processing (e.g. the cognitive control of emotion).

As expected, spatiotemporal neural clusters involving stimulation and rational choice and feedback effects occurred in mid-latency or late time intervals (earliest effect starts at 220ms). This is probably due to the nature of rational processing, as a higher-order cognitive process, that is connected downstream to sensory stimulus processing and emotionally driven decisions. ERP results confirm this pattern, where correlates of rational decision-making are typically not found earlier than the P300 (Wichary et al., 2017). In comparison, clusters that are more related to impulsive decision-making have their onset already in rather early time ranges (Fig. SM3, SM4, SM5A).

Despite these new insights in vmPFC-functioning and non-invasive brain stimulation in decision-making and feedback-processing, there are limitations to consider. First, to guarantee successful blinding of participants to the stimulation conditions and to reduce inter individual variance we opted here for a within-subjects-design and first sacrificed the comparison with a sham condition (i.e. participants easily detect the difference between active and sham tDCS while both active conditions are typically undistinguishable; see SM2.1). While all conclusions regarding causal functionality and modulating capability of the vmPFC remain unaffected, it remains to be resolved whether inhibitory stimulation might solely evoke temporary vmPFC dysfunctions in healthy controls as reported in patients and/or if excitatory stimulation might improve rational decision-making even in healthy participants. To investigate this question, follow-up studies should use between-designs comparing effects of excitatory versus sham vmPFC-tDCS in healthy participants. Such designs could also be used in clinical settings to test a complementary treatment option for gambling (and other behavioral) addictions. In fact, first indications for positive effects of mPFC stimulation have been demonstrated for obsessive-compulsive disorders (Adams et al., 2021), which is likewise associated with impaired impulse-control. Furthermore, the neural data of the feedback phase contained limited informative value, due to the extremely strong differential effects of the outcomes (loss >> gain, driven by loss-aversion), which explained most variance and potentially masked relevant interaction effects. Nevertheless, we could at least partly circumvent this problem by calculating a t-test, that was justified by our hypotheses and by the behavioral effects.

## Conclusion

Our results not only support the claim that the vmPFC plays a causal role for rational decision-making and feedback-processing, but they also indicate that rationality can be modulated by its non-invasive stimulation. Improved decision-making after excitatory vmPFC-tDCS as reflected in maximized wins and decreased susceptibility to the framing-effect is a compelling illustration of this influence. The same is true for the feedback phase, where less loss-biased stimulus processing might guide more adaptive behavior in the future. These higher-order cognitive processes were reflected in prefrontal clusters of neural activity at mid-latency to late time intervals. They putatively indicate that loss-aversion can be attenuated leading to more rational decision-making. The finding that vmPFC-excitation compared to inhibition can induce more rational decision-making and feedback-processing raises the hope for clinical applications of vmPFC-tDCS in addiction disorders such as pathological gambling.

## Methods

### Participants

We included 37 (17 women) right-handed volunteers, aged 19 to 29 years (*M* = 23.42, *SD* = 2.70) meeting the inclusion criteria (see SM1.1). Participants were pseudo-randomly assigned to experimental groups that were matched regarding demographic and psychometric features. The study was ethically approved by the ethics committee of the medical school at the University of Münster.

Participants were told a cover story to ensure authentic gambling behavior. It was stated that they could win an amount between 0 and 36$ in addition to the fixed allowance of 30$. Following the study, participants were elucidated regarding the cover story and everybody received the full amount of 66$.

### Specifications on the Gambling Task

Each trial began with a fixation cross after which the initial amount of money was presented, for which the participants could successively gamble with a specified risk (game stake of 25ct, 50ct, 75ct or 100ct). Second, the ‘choice stimulus’ appeared on which basis the participants had to decide whether they would choose a safe (‘keep’) or a risky (‘gamble’) option. The framing-effect is relevant in the ‘keep’ option: If the game stake was for instance 50ct (Fig 1A), the green gain-frame informed about a safe win of 20ct (i.e. equivalent to a safe loss of 30ct) while the red loss-frame predicted a safe loss of 30ct (i.e. equivalent to a safe win of 20ct). According to this scheme the gain-frames informed about safe wins of 10ct, 30ct, or 40ct and the loss-frames of safe losses of 15ct, 45ct, or 60ct, if the initial amounts were 25ct, 75ct or 100ct, respectively.

### Experimental procedure

In a within-subjects-design each participant received excitatory/anodal and inhibitory/cathodal stimulation over the course of two sessions with a minimum-interval of 48 hours between the sessions (Methods Fig. 1). The allocation of stimulation order (excitatory or inhibitory stimulation first) was randomized across participants. In the beginning of the first session, participants gave written informed consent, filled questionnaires, comprising the Beck Depression Inventory (Beck et al., 1996), the Reward Responsiveness scale (Van den Berg et al., 2010), the Intolerance of Uncertainty scale (Gerlach et al., 2008), and the Social Desirability Scale (Crowne and Marlowe, 1960). After tDC stimulation, participants executed the gambling task in the MEG, where event-related fields (ERFs) in response to the choice- and feedback stimuli were measured. At the end of each session, participants rated the feedback regarding subjective hedonic valence and emotional arousal on a self-assessment manikin (SAM) rating scale (Bradley and Lang, 1994), rated their state mood on the Positive and Negative Affect Schedule (PANAS; Watson, Clark &Tellegen, 1988) and rated their perceived stimulation pleasantness and stimulation intensity on an in-house questionnaire. In the second session, the same procedure was used with the opposite stimulation polarity. Finally, subjects were elucidated about the cover story. The duration of both sessions combined was approximately 200 minutes.

**Methods Figure 1.**
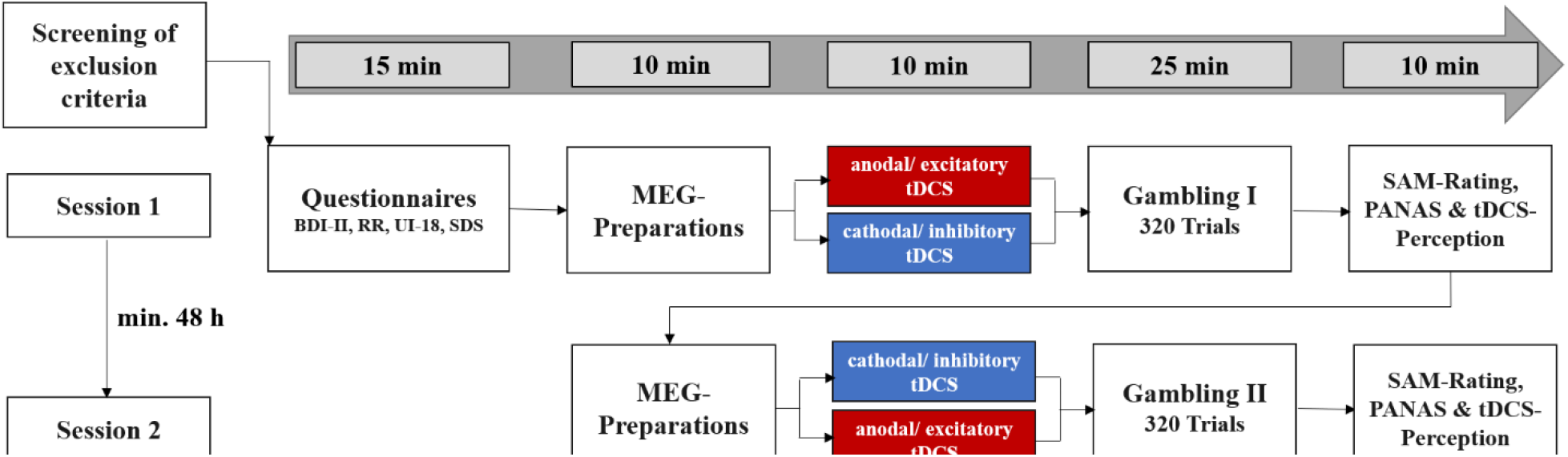
Overview over the experimental procedure. *Abbreviations:* BDI-II: Beck Depression Inventory-II. RR: Scale for Measuring Reward Responsiveness. UI-18: Intolerance of Uncertainty scale. SDS-CM: Social Desirability Scale by Crowne and Marlowe. SAM-Rating: Subjective Ratings of Hedonic Valence and Emotional Arousal. PANAS: Positive and Negative Affect Schedule. For results of the questionnaires see SM1.1.

### tDCS

Transcranial direct current stimulation (tDCS) is a widespread and effective method to modulate brain activity from outside the skull. An anodal or excitatory stimulation depolarizes the membrane potential of neurons, what increases their excitability depending on the strength of the applied electric field. In contrast, cathodal or inhibitory stimulation hyperpolarizes the neuron membrane, reducing the likelihood of action potentials (Sparing and Mottaghy, 2008). An important advantage of tDCS is the low rate of side effects (e.g. headache, nausea and insomnia) and, with particular relevance for ventral prefrontal target regions, the absence of unwanted co-stimulation of facial and ocular muscles and nerves. Changes of cortical excitability can last up to one hour after a single stimulation (Poreisz et al., 2007). We implemented the tDCS-montage as used in our previous studies to non-invasively stimulate the vmPFC (Junghofer et al., 2017; Roesmann et al., 2021; Winker et al., 2020, 2019, 2018). The active electrode was placed on the forehead (3 × 3 cm) and the extracephalic reference under the chin (5 × 5 cm). Electrodes were plugged into sponges that were soaked with a sodium-chloride solution to ensure electric conductivity. For excitatory/anodal or inhibitory/cathodal stimulation, the forehead electrode was used as anode or as cathode respectively. This electrode-montage results in maximal stimulation of the vmPFC and minimal stimulation of adjacent brain regions, as revealed by finite-element based forward modelling of tDCS currents (Wagner et al., 2014). Using a DC Stimulator Plus (NeuroConn GmbH), we applied a maximum current of 1.5 mA for 10 minutes with both stimulation polarities.

**Methods Figure 2.**
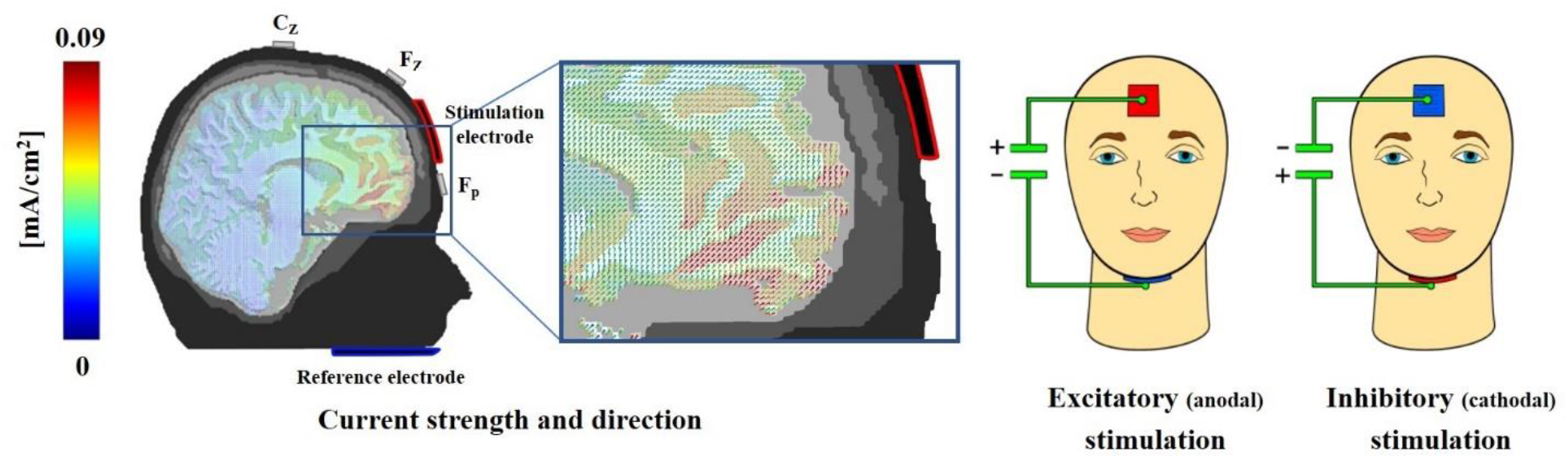
An iterative gain function algorithm aiming at maximal vmPFC stimulation revealed an electrode positioning with a small mid-frontal electrode and an expanded extracephalic chin reference. This array allowed a quasi-reference-free stimulation, providing clear differentiation of excitatory and inhibitory effects. Participants were stimulated at two different days for 10 min with 1.5 mA in an either excitatory (anodal forehead electrode) or inhibitory (cathodal forehead electrode) fashion. While the current strength is identical for anodal and cathodal stimulations, the direction of effects, as indicated by cones in the magnification, is reversed. A modeled 1.5mA stimulation resulted in a maximum current density in the vmPFC regions of approximately 0.09 mA/cm2 (red colors). Actually, all sponges had the same color to prevent any inferring of the participants based on sponge color. This figure was published first in Junghoefer et al. *Cerebral Cortex* (2017).

### Recording and preprocessing of MEG

Event-related fields were measured using a 275 whole-head sensor system (CTF Systems, first-order axial gradiometers) with a sampling rate of 600 Hz across a frequency range from 0 to 150Hz (anti-aliasing hardware filtering). The continuous data were down-sampled to 300Hz and filtered with a 0.1 high-pass-filter and 48 Hz low-pass-filter. We extracted epochs from 200ms before and 600ms after stimulus onset and employed the interval of -150ms to 0ms for baseline adjustment. To identify and reject artifacts the method suggested by Junghöfer and colleagues was used (Junghöfer et al., 2000). With this method, individual and global artifacts were discovered. In case noisy channels were identified, their signal was estimated by spherical-spline-interpolation based on the weighted signal of all remaining sensors. A minimum threshold of 0.01 for the estimated Goodness of Interpolation was applied and trials exceeding this value were rejected. If more than 30% of the trials were discarded in any session, the respective participant was excluded from further analysis (three participants). Trials within each experimental condition were averaged for each participant and session and the underlying neural sources of the measured ERFs were estimated by applying L2-Minimum-Norm-Estimates (L2-MNE; Hämäläinen &Ilmoniemi, 1994). The preprocessing and analysis of the MEG data was performed with the MATLAB (2019b)-based EMEGS software (version 3.1; Peyk, De Cesarei &Junghöfer, 2011). For details see SM1.2.

### Data-Analysis

We used mixed effects models for the behavioral data because of their robustness in repeated-measures compared to conventional models (Baayen et al., 2008). Detected effects were then resolved with traditional post-hoc tests.

Aiming to test which factors influence rational decision-making (choice to ‘keep’ or ‘gamble’) we calculated a mixed effects logistic regression with the predictors stimulation (excitatory, inhibitory), risk-to-lose (20%, 40%, 60%, 80%) and frame (gain-frame, loss-frame). Since we were mainly interested in the interaction effects of stimulation by frame and stimulation by risk-to-lose we modeled random effects for these interactions. The overall model was significant (*χ*²(16) = 13297.00, *p* < 0.001) and within this model all main effects got significant: stimulation (*z* = -9.69, *p* < 0.001), risk-to-lose (*z* = -9.60, *p* < 0.001) and frame (*z* = 2.64, *p* = 0.008). The interaction of stimulation by frame was insignificant (*z* = 0.39, *p* = 0.695), but since we had specific hypothesis regarding this effect, we calculated a separate logistic regression with only these predictors. As expected, the interaction of stimulation by frame was significant in this model (*z* = 2.19, *p* = 0.029). However, the interaction of stimulation by risk-to-lose got highly significant in the overall model (*z* = 6.60, *p* < 0.001).

The neural analyses in the decision-making phase (stimulation by frame by decision and stimulation by risk-to-lose) were performed separately from each other to ensure a sufficient number of trials per condition for a reasonable signal-to-noise-ratio in the source estimation. To correct for multiple comparisons we applied a non-parametric approach proposed by Maris and Oostenveld (Maris and Oostenveld, 2007). For details see SM1.4.For the analysis of the perceived hedonic valence (SAM-rating; Bradley &Lang, 1994), 9-point Likert scale) of the feedback, we computed a mixed effects linear regression with the predictors stimulation (excitatory, inhibitory), outcome (gain, loss) and decision (keep, gamble). Since we were mainly interested in the interaction effects of stimulation by decision, we modeled random effects for this interaction.

The overall model was significant (*χ*²(9) = 526.44, *p* < 0.001) and revealed significant main effects of stimulation (*t* = -2.99, *p* = 0.003), outcome (*t* = -25.08, *p* < 0.001) and decision (*t* = -5.16, *p* < 0.001). While the two-way-interactions of stimulation by decision (*t* = -0.17, *p* = 0.867) and stimulation by outcome (*t* = 1.65, *p* = 0.245) were insignificant, the three-way-interaction got highly significant (outcome by decision by stimulation: *t* = 5.33, *p* < 0.001).

