## Supplementary Material for "Noninvasive stimulation of the ventromedial prefrontal cortex modulates rationality of human decision-making"

### SM1. Methods

#### 1.1. Participants

The recruitment of participants was conducted via social media and the participant pool of our institute. Exclusion criteria were current or lifetime psychiatric diagnosis, psychopharmacological treatment, current or past psychotherapy, neurological or severe somatic disease, prior participation in a gambling study and pregnancy. One participant was excluded due to a misunderstanding of the experimental paradigm.

BDI-II scores were measured to exclude participants with even moderate signs of depression to prevent potentially adverse effects of the inhibitory vmPFC stimulation on mood. No participant had to be excluded from the study due to this criterion.

The reward responsiveness scale (RR; Van den Berg et al., 2010) and the Intolerance of Uncertainty scale (UI-18; Gerlach et al., 2008) were measured to potentially exclude participants with extremely low risk tolerance, while the social desirability scale (SDS; Crowne and Marlowe, 1960) was evaluated to potentially exclude participants who were extremely concerned with social approval. No outlying participants were observed on these scales.

**Table SM1**

Demographic and psychometric characteristics of participants

| | **M** | **SD** | $\chi^2$ | **df** | **p** |
|---|---|---|---|---|---|
| **Demographic Characteristic** | | | | | |
| N | 37 | | | | |
| Female (N, %) | 17 (45.9) | - | *0.62* | 1 | 0.243 |
| Stimulation Order (Exc-Inh, %) | 19 (51.4) | - | *0.03* | 1 | 0.869 |
| Age (years) | 23.42 | 2.70 | | | |

| | M | SD | $\chi^2$ | df | p |
|---|---|---|---|---|---|
| **Psychometric Characteristics** | | | | | |
| BDI-II | 2.78 | 2.65 | | | |
| RR | 25.70 | 2.65 | | | |
| UI-18 | 40.60 | 10.78 | | | |
| SDS | 14.02 | 3.64 | | | |
| PANAS-Pos-Exc | 30.08 | 6.63 | | | |
| PANAS-Pos-Inh | 29.83 | 6.80 | | | |
| PANAS-Neg-Exc | 11.61 | 2.35 | | | |
| PANAS-Neg-Inh | 12.41 | 4.16 | | | |

*Note.* Stimulation order – excitatory first, inhibitory second: N = 21; inhibitory first, excitatory second: N = 20. BDI-II = Beck Depression Inventory (Beck et al., 1996). RR = Reward responsiveness scale (Van den Berg et al., 2010). UI-18 = Intolerance of Uncertainty scale (Gerlach et al., 2008). SDS-CM = Social desirability scale (Crowne and Marlowe, 1960). PANAS = Positive and negative affective schedule (Watson et al., 1988).

#### 1.2. Recording and preprocessing of MEG

We conducted MEG measurements with a 275 whole-head sensor system (CTF Systems) with first-order axial gradiometers to record the visually evoked magnetic fields (Omega 275; CTF, VSM MedTech Ltd., Coquitlam, Canada). Frequencies between 0 and 150 Hz were gathered with a sampling rate of 600 Hz. To capture the participants individual head shapes a 3D tracking device (Polhemus, Colchester, VT, USA; http://www.polhemus.com/) was used and the position in the scanner was ascertained by three landmark coils in the ears and on the nasion.

Experimental conditions of the choice stimulus (decision-making) contained the factors game stakes (25ct, 50ct, 75ct and 100ct), risk-to-lose condition (20%, 40%, 60%, 80%), decision (gamble, keep), frame (gain frame, loss frame) and stimulation (excitatory, inhibitory). Since the accuracy of inverse source modeling (see below) does strongly depend on the signal-to-noise ratio, which increases with the square-root of the number of trials averaged per condition, we averaged across factors that were not relevant for a distinct research question. Testing the impact of stimulation and framing on decision (see Fig.1C main text), we thus averaged across the factors game stakes and risk-to-lose condition. Testing for the impact of stimulation on the factor risk-to-lose condition (see Fig.2D main text), we averaged across the factors game stakes, decision and frame. Experimental conditions of the feedback stimulus (feedback processing) contained the factors game stakes, risk-to-lose condition, decision, frame, stimulation and outcome (gain, loss). Testing the impact of stimulation on outcome in the relevant 'keep' condition (see Fig. 3C main text), we averaged across the factors game stakes, risk-to-lose condition and decision.

Following the averaging, the underlying neural sources were estimated by the L2 minimum-norm estimation (Hämäläinen and Ilmoniemi, 1994), an inverse modeling technique that does not need *a priori* assumptions about the location and distribution of neural generators. A spherical model with 350 evenly distributed dipole pairs (azimuthal and polar direction) and a source shell radius approximately corresponding to the grey matter depth – i.e., 87% of the individually fitted head – was employed as a source model. The Tikhonov regularization parameter lambda was set to 0.1. The source-direction-independent neural activities (vector length of the estimated source activities at each position) were estimated for each individual participant, condition and time point. In addition to the three participants excluded during artifact inspection, further participants were excluded due to outlier analyses conducted in source space, which were conducted for each of the three neural analyses separately. Regarding the analysis stimulation by decision by frame (decision-making), three participants had to be excluded (final sample: N = 30). In the analysis stimulation by risk-to-lose condition (decision-making), four participants had to be excluded (final sample: N = 29). Finally, in the analysis stimulation by decision by outcome (feedback processing), five participants had to be excluded (final sample: N = 28). In these subjects, in one or both sessions (inhibitory or excitatory stimulation), either the mean of the standard deviation between experimental conditions across time or the maximum of normed (by mean) standard deviation between experimental conditions differed from the sample median by more than four standard deviations. This resulted in the smallest shared sample of 25 participants for the MEG analysis. All behavioral and neural analyses were conducted again with this smallest shared sample, and all reported results remained qualitatively equivalent. All effects also remained qualitatively equivalent when stimulation order (excitatory or inhibitory first) was employed as additional factor.

##### 1.3. Analysis of the behavioral data

Behavioral measures were analyzed using the statistics program R, applying an α-level of 0.05. If suitable, Greenhouse-Geiser corrected statistics were reported.

##### 1.4. Analysis of neural data

We applied intervals of interest from 0 – 300ms to investigate early bottom-up processes and 300 – 600ms for later more cognitive processes, as we did in previous studies (Roesmann et al., 2021; Winker et al., 2018). If the temporal extent of any resulting spatio-temporal cluster reached the predefined border of 300ms, the interval was stepwise extended by 50ms. To tackle the problem of multiple comparisons, we used a non-parametric correction approach (Maris and Oostenveld, 2007). In this procedure, statistical values (i.e., $F$-values of the respective ANOVA analyses, t values of t-tests, r values for correlational analyses) per time point and estimated test dipole entered so-called spatio-temporal cluster masses if the respective test statistic exceeded a critical alpha level of $p = 0.01$ (sensor-level criterion). The cluster mass was calculated as the three-dimensional spatio-temporal integral of the statistical values (i.e., the statistical value reflects the 'density' within the cluster volume, resulting in the 'mass' of the cluster). In a second step, cluster masses within the pre-defined intervals of interest (early: 0-300ms, late: 300-600ms) and within the prefrontal region of interest (based on anterior vs. posterior characterization of the head model implemented in EMEGS) were tested against cluster masses of identical test statistics, yielded by 1,000 permuted drawings from the same dataset. The distribution of the permutations was employed to determine the cluster criterion. If the actual cluster mass exceeded this critical cluster mass of $p = 0.05$ (i.e., greater than 95% of the biggest cluster masses of each of the 1,000 permuted drawings; cluster-level criterion), the cluster was classified as significant. The duration of the clusters was rounded to the nearest 10ms, since this reduces the risk of excessive accuracy for the given temporal resolution of cluster onsets and offsets. This procedure was implemented for all examined effects.

For visualization purposes, spherical L2-MNE topographies were projected on standard 3D brain models.

### SM2.Results and Discussion

#### 2.1. tDCS perception

No significant differences between excitatory and inhibitory stimulation in perceived hedonic valence ($t(35) = -1.27$, $p = 0.212$) and intensity ($t(35) = -0.13$, $p = 0.900$) emerged. Thus, participants were not able to differentiate between the stimulation conditions.

#### 2.2. Investigation of mood effects

To control for modulations of mood effects by the stimulation, we calculated a 2x2 repeated-measures ANOVA with the factors stimulation (excitatory, inhibitory) and affect (positive PANAS scale, negative PANAS scale). The analysis did not reveal a main effect of stimulation ($F(1,35) = 0.92$, $p = 0.527$) nor an interaction of affect by stimulation ($F(1,35) = 0.47$, $p = 0.498$).

##### 2.3. Supplementary analyses on decision-making

###### 2.3.1. Reaction times

A linear regression revealed a slight but significant influence of stimulation on reaction times ($F(1, 23038) = 16.616$, $p < 0.001$). After excitatory tDCS ($M = 1.27$ sec), reaction times were prolonged compared to inhibitory stimulation ($M = 1.20$ sec), which was particularly pronounced in the early and middle phase of the gambling paradigm (see Fig. SM1).

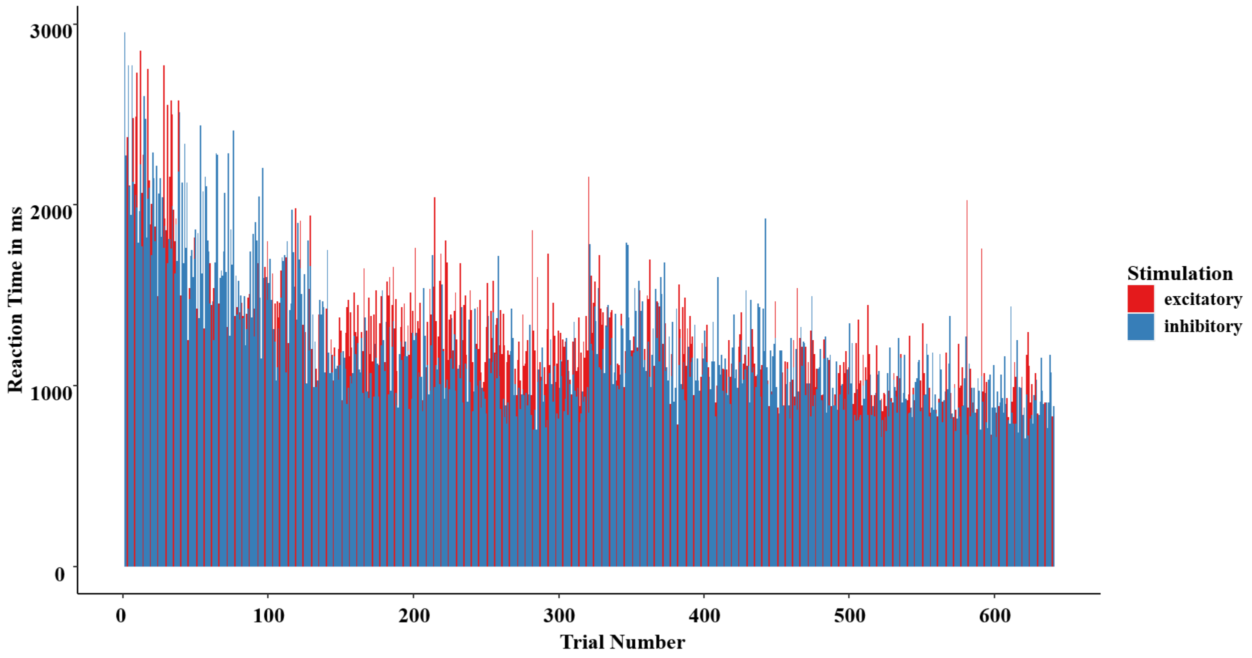


*Figure SM1.* Reaction times (decision to ‘keep’ or ‘gamble’) in dependency of trial number and stimulation condition. Following excitatory stimulation, reaction times were slightly prolonged compared to inhibitory stimulation. This effect particularly occurred in the early and middle phases of the gambling paradigm and might, thus, reflect less impulsive decision-making after vmPFC excitation or more impulsive decisions after inhibitory stimulation.

In conjunction with the behavioral and neural results reported in the main text this could be interpreted as further sign for less impulsive decision-making after vmPFC-excitation or more impulsive decisions after inhibitory stimulation. This is also in line with the time course of the neural results, since clusters that implied greater impulsivity occurred at rather early latencies (Fig. SM4A and SM5), while effects that were related to increased rationality emerged with later onset (Fig. 1C and 2C in the main text and Fig. SM5B). The fact that effects of stimulation on reaction times were more pronounced in early and middle stages of the gambling paradigm could reflect an expected logarithmic reduction of the impact of the 10 minutes of tDCS stimulation across the subsequent 25 minutes of gambling. It might however also suggest a formation of more elaborate strategies after excitatory stimulation at the beginning of the gambling task, which were then used for the ongoing paradigm.

**2.3.2. Interaction of stimulation and game stake**

To strengthen the participants' feeling of being in a real gambling situation and reduce the participants' probability of calculating the expected values (ecological validity), the game stakes varied in each trial between 25ct, 50ct, 75ct and 100ct. This variation, however, also allowed us to investigate how responsibly participants gambled depending on stimulation. Therefore, we calculated a logistic regression on the interaction of stimulation (excitatory, inhibitory) by initial amount (25ct, 50ct, 75ct, 100ct), which became significant ($z = 2.55$, p = 0.011; see Fig. SM2). With increasing game stakes, risk-taking behavior increased after vmPFC inhibition (post-hoc analysis inhibitory only: $z = 4.25$, $p < 0.001$), while risk-taking behavior was not modulated by the initial amount after excitatory stimulation (post-hoc analysis excitatory only: $z = 0.65$, $p = 0.515$).

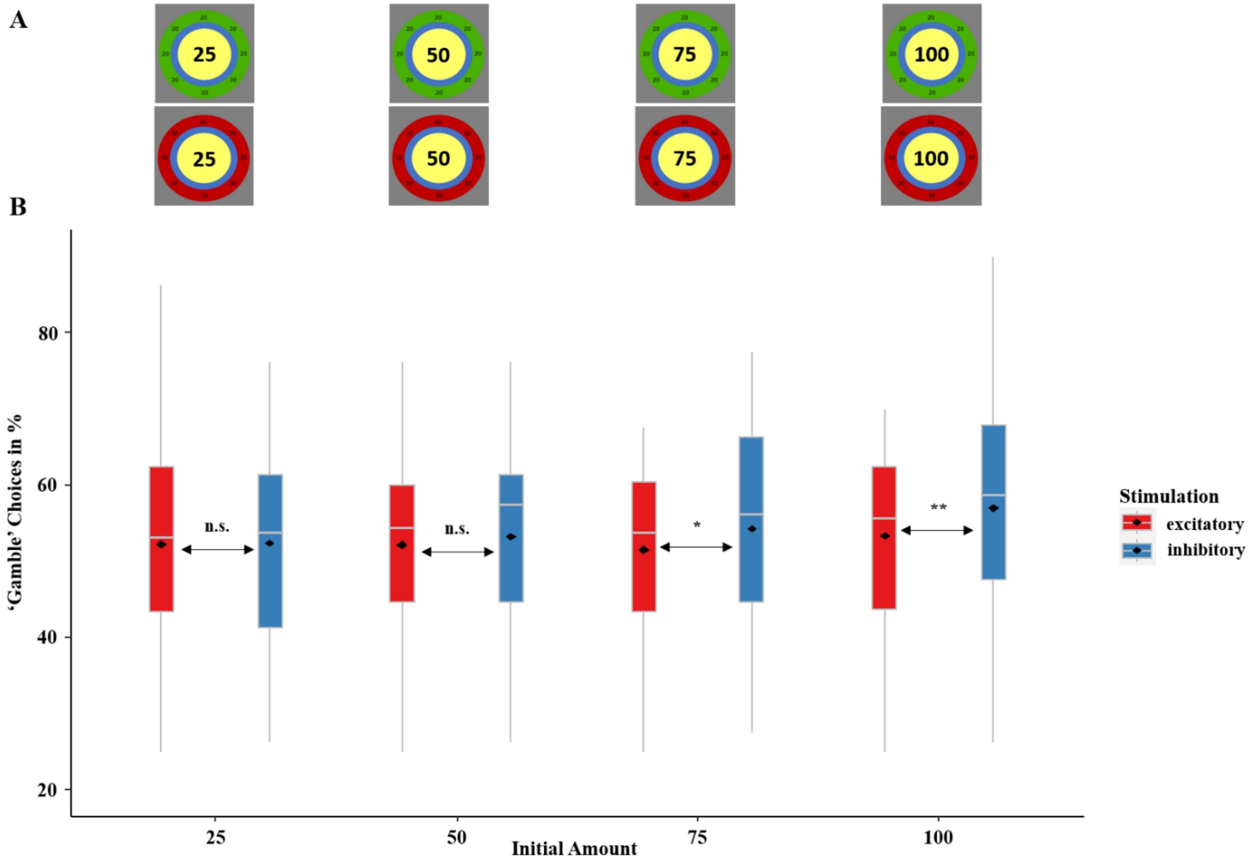


*Figure SM2*. **A.** Toward increased ecological validity, the game stakes were varied in each trial between 25ct, 50ct, 75ct and 100ct. **B.** Proportion of 'gamble' choices depending on the initial amount (game stakes) and stimulation in percentage. After inhibitory stimulation, participants showed an increasing tendency to gamble with increasing game stakes, while gambling behavior was not affected by the initial amount after excitatory stimulation. Boxplots indicate means (black dots), medians (grey lines) and lower and upper quartiles. Asterisks indicate significance levels: + < 0.1, * < 0.05, ** < 0.01, *** < 0.001.

This could be interpreted as a further sign that excitatory vmPFC-tDCS might help to restore more normal gambling behavior, since gambling addicts exhibit vmPFC hypoactivation, and patients with vmPFC lesions tend to risk greater amounts to equalize previous losses (Reuter et al., 2005; Xue et al., 2011), while healthy controls are able to disengage from risky behavior after multiple losses (Shiv et al., 2005). Or, put the other way around, inhibitory vmPFC-tDCS appears to lead to increased susceptibility for uneconomical gambling behavior.

##### 2.3.3. Neural main effects

The 2x2x2 ANOVA revealed a significant main effect of frame at right anterior temporal and orbitofrontal areas ranging from 70-170ms (*p*-cluster = 0.031). Neural activity in this cluster in response to the 'gain-frame' was stronger than responses to the 'loss-frame' ($F(1, 29) = 9.06$, $p = 0.005$, see Fig. SM3). This cluster spatially overlaps but precedes the subsequent interaction of frame by stimulation (220-270ms; Fig. 1C main text). The later frame by stimulation interaction has been interpreted as reflecting a relatively reduced inhibition of loss-aversion after excitatory stimulation and relatively enhanced inhibition after inhibitory stimulation.

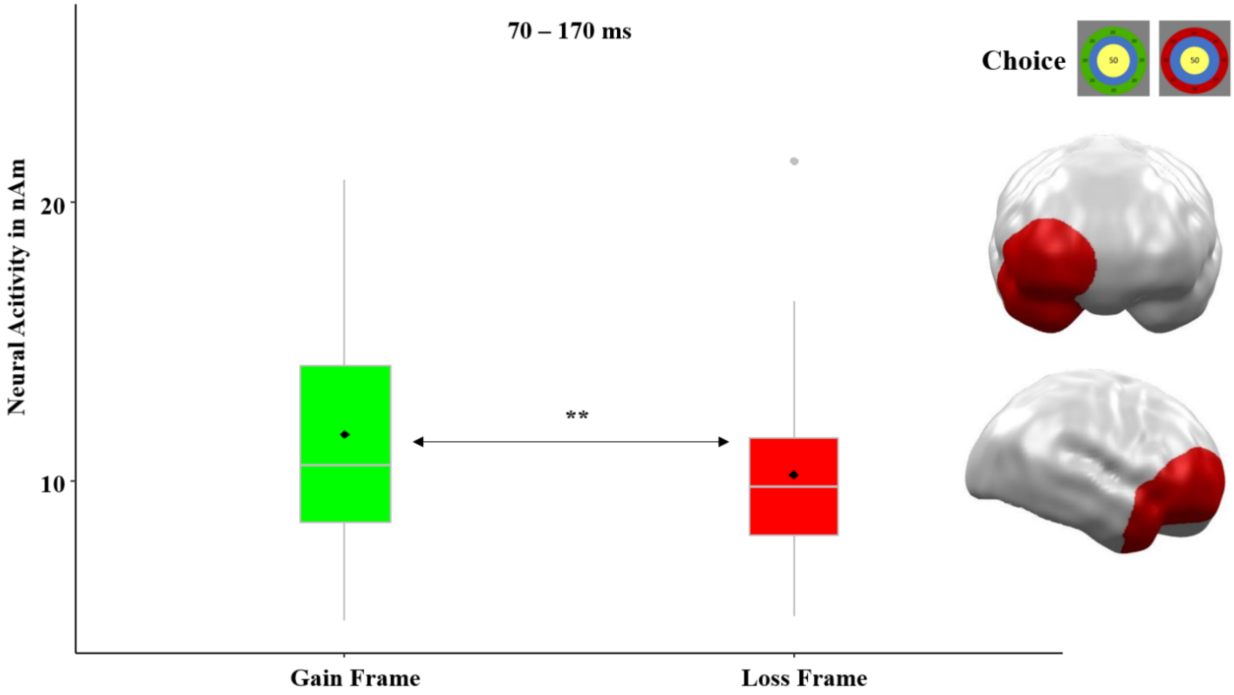

*Figure SM3.* Significant prefrontal spatio-temporal cluster featuring a main effect for frame (gain, loss) by an ANOVA. Topographies of effects observed in L2-MNE were projected on standard 3D brain models for visualization. Boxplots indicate means (black dots), medians (grey lines) and lower and upper quartiles. Asterisks indicate significance levels: + < 0.1, * < 0.05, ** < 0.01, *** < 0.001.

This earlier main effect of frame could likewise be interpreted as an early correlate of loss aversion, with relatively reduced inhibition of potential losses compared to potential gains (i.e., enhanced 'gating' of losses) eventually leading to the framing effect. It remains to be resolved in future studies whether the chosen tDCS stimulation was not sufficiently strong (at 1.5 mA) or long (at 10 minutes) to prove a modulation of the frame effect also in this earlier time interval or whether the vmPFC can modulate only later stages of processing in this region.

Additionally, we calculated paired t-tests to detect main effects of stimulation (excitatory, inhibitory) and decision (keep, gamble). The t-test for stimulation revealed a significant main effect ($p$-cluster = 0.013) spanning effectively the whole analyzed time interval (10 – 600ms). As expected, observed neural activations at the vmPFC region of stimulation and neighboring areas were enhanced after excitatory compared to after inhibitory stimulation ($t(29) = 2.38$, $p = 0.024$, see Fig. SM4).

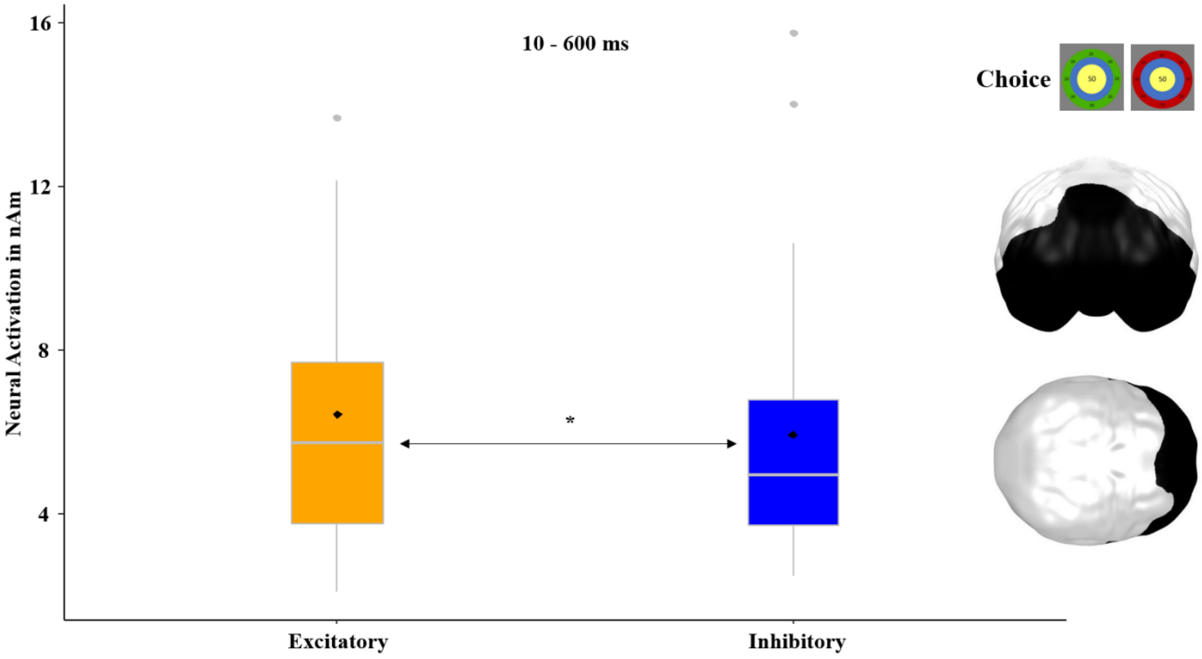


*Figure SM4.* Significant prefrontal spatio-temporal cluster featuring a main effect of stimulation as revealed by a t-test. Topographies of effects observed in L2-MNE were projected on standard 3D brain models for visualization Boxplots indicate means (black dot), medians (grey line) and lower and upper quartiles. Asterisks indicate significance levels: + < 0.1, * < 0.05, ** < 0.01, *** < 0.001.

This finding can be interpreted as proof for the intended experimental manipulation (see Methods Fig. 2 main text). The corresponding analysis for a main effect of stimulation in the feedback phase remained insignificant, probably due to the strong effects of the outcome (loss >> gain), which might have masked other effects by explaining most of the variance.

The t-test for the main effect of decision revealed two significant clusters. A first early to mid-latency effect (10[1] – 340ms) emerged at left dorsolateral and dorsomedial prefrontal areas (*p*-cluster = 0.010) and featured greater activity for later ‘gamble’ choices in comparison to ‘keep’ choices ($t(29) = -2.49$, $p = 0.019$; Fig. SM5A). This cluster overlaps temporally and spatially with the dorsal part of the cluster, which showed increasing neural activity with increasing cumulative wins (i.e., with increasing rational choice behavior; see Fig. SM8) for the choice stimulus across participants. Since the ‘gamble’ option was more rational than the ‘keep’ option in the majority of choices (see main text Fig. 2B), this stronger neural activation for ‘gamble’ might again reflect relatively increased necessary effort for rational choice behavior (e.g., to suppress loss aversion). Further support for this assumption comes from an fMRI study that used a very similar paradigm (Rogers et al., 2004). Just as our MEG cluster indicates, greater blood-oxygen levels were reported from Rogers and coworkers in the left hemisphere when the ‘gamble’ option was chosen in the decision phase. As for the above main effect of frame (Fig. SM3), it remains to be resolved whether the chosen stimulation had sufficient impact to prove a modulation on decisions also in this earlier time interval or whether the vmPFC modulates only later stages of processing in this dorsal region.

[1] It should be noted that to prevent latency shifts, MEG data has been filtered in a forward-backward fashion. As backward filtering smears effects to time points preceding the real effects, statistical effects might occur even preceding the real onset of effects. Thus, the ‘real’ onset of this effect might occur later than 10ms.

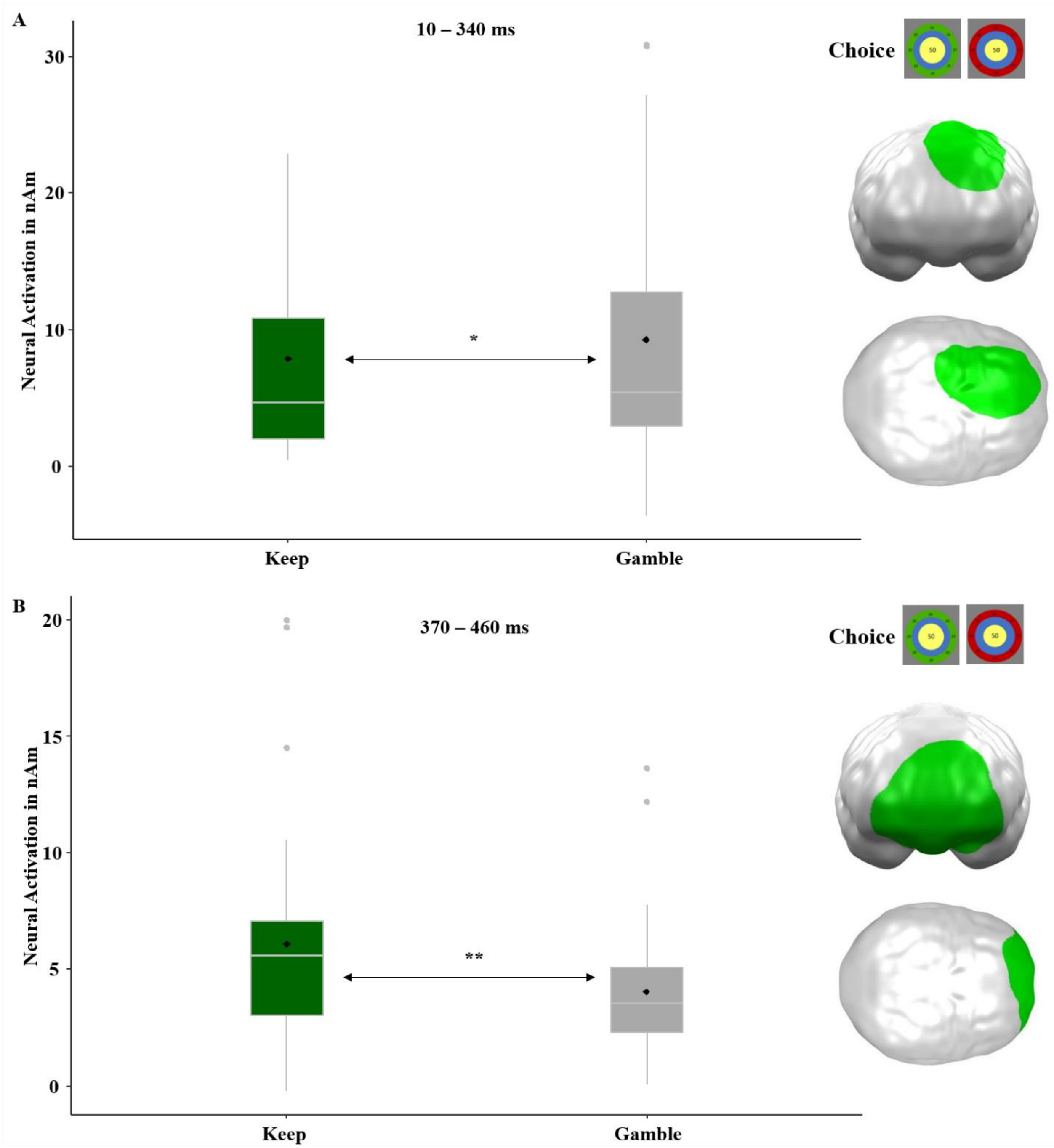


*Figure SM5.* Significant spatio-temporal clusters featuring a main effect for decision revealed by a t-test. **A.** Left and medial prefrontal brain regions with greater activations when the ‘gamble’ option was chosen. **B.** Medial prefrontal brain regions with greater activations when the ‘keep’ option was chosen. Topographies of effects observed in L2-MNE were projected on standard 3D brain models for visualization Boxplots indicate means (black dot), medians (grey line) and lower and upper quartiles. Asterisks indicate significance levels: + < 0.1, * < 0.05, ** < 0.01, *** < 0.001.

A second cluster occurred in a subsequent time interval at the area of stimulation at ventromedial prefrontal regions (370-460ms; $p$-cluster = 0.005; Fig. SM5B), showing an opposite pattern with stronger activation for a later ‘keep’ compared to a later ‘gamble’ decision (keep > gamble; $t(29) = 3.03$, $p = 0.005$). As this region typically shows enhanced activity for positively evaluated stimuli (Sescousse et al., 2013) and reduced activity for negatively evaluated ones (Lindquist et al., 2016), enhanced neural activity in this areas should have been expected in response to stimuli that were evaluated pleasant enough to trigger a ‘keep’ decision, while relatively reduced activity in this region (more negative evaluation) should have preferentially triggered to the alternative ‘gamble’ option. It should be noted that this effect of decision, although occurring at the target region of stimulation, was not modulated by the stimulation. Based on our hypothesis that vmPFC-tDCS should modulate valence biases, an interaction with modulation could have been expected in the vmPFC region. This might be consequent upon the rather strong main effect of stimulation in this region (see Fig. SM4), which might have covered smaller interaction effects. However, it also matches with previous studies in our lab, in which interactions with the factor stimulation occurred in remote areas but not in the vmPFC target area of stimulation (Junghofer et al., 2017; Winker et al., 2020, 2019, 2018).

##### 2.3.4. Neural interaction effects

The ANOVA revealed two further clusters with significant interactions with the factor stimulation, that were not reported in the main text because the interaction either occurred in a very early time interval where effects on rational decision-making has not been predicted (Fig. SM6) or because it revealed an unpredicted three-way interaction (Fig. SM7), which asks for replication in future studies.

First, an interaction effect of stimulation by frame appeared in a very early time interval from 0[2] to 80ms in superior medial prefrontal areas (*p*-cluster = 0.021). Here, in the gain-frame, excitatory stimulation resulted in stronger neural activity compared to inhibitory stimulation while the opposite pattern occurred in the loss-frame condition ($F(1,29) = 13.89$, $p = 0.001$).

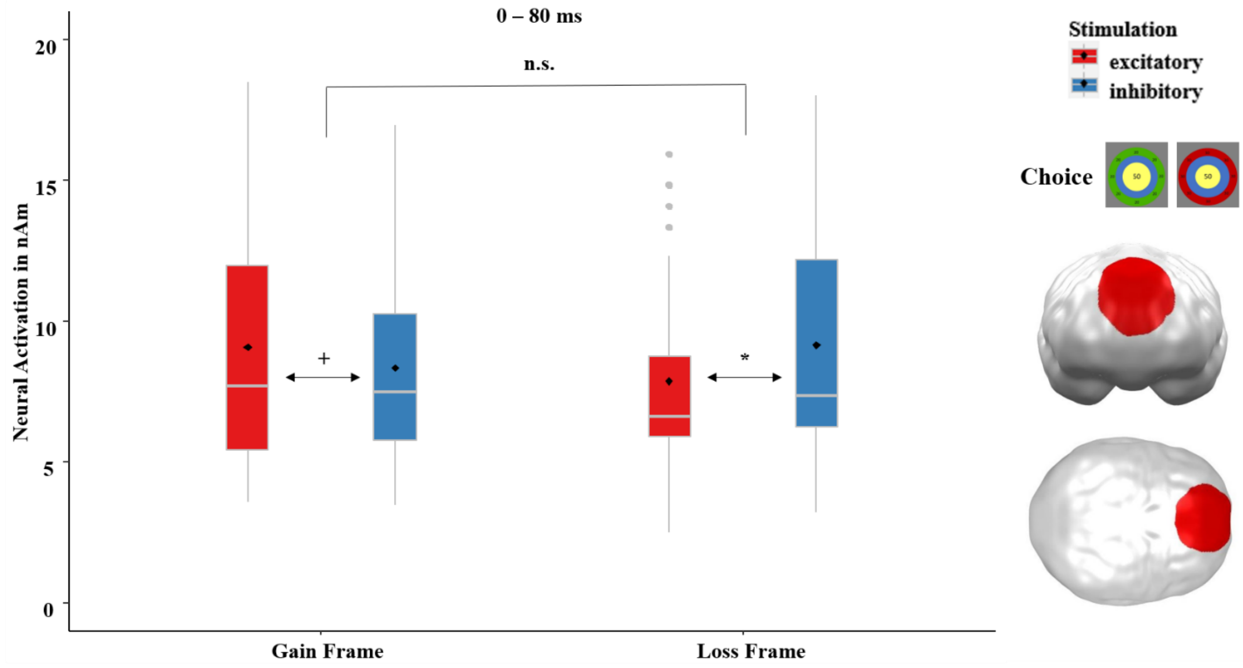


*Figure SM6.* Significant medial prefrontal spatio-temporal cluster featuring an interaction effect of stimulation by frame revealed by an ANOVA. Topographies of effects observed in L2-MNE were projected on standard 3D brain models for visualization. Boxplots indicate means (black dots), medians (grey lines) and lower and upper quartiles. Asterisks indicate significance levels: + < 0.1, * < 0.05, ** < 0.01, *** < 0.001.

This cluster directly precedes and nicely overlaps with the dorsal part of the cluster (Fig. SM8), which showed increasing neural activity with increasing cumulative wins for the choice stimulus across participants (although this correlation effect revealed no significant impact of stimulation). It might thus very well be interpreted as reflecting a reduced loss processing and increased gain processing after excitatory compared to after inhibitory stimulation, eventually

---

[2] It should be noted that to prevent latency shifts, MEG data has been filtered in a forward-backward fashion. As backward filtering smears effects to time points preceding the real effects, statistical effects might occur even preceding the real onset of effects. Thus, the 'real' onset of this effect might reasonably occur after stimulus onset.

leading to more rational choice behavior, which would match our hypothesis. However, this cluster (Fig. SM6) also overlaps with the cluster which showed an increased activation in response to loss frames in the ‘keep’ option after excitatory compared to inhibitory stimulation (see Fig. 3C main text). This effect has been interpreted as increased inhibition of loss feedback after excitatory stimulation. Though this effect (Fig. SM6) occurred at choice processing and not at feedback processing (as Fig. 3C main text) and occurred at very early (0-80ms) and not at late time intervals (470-510ms), we cannot ignore that an interpretation as reduced loss inhibition after excitatory stimulation would be contrary to our hypotheses. Alternatively, such an inverted effect in a very early time interval might reflect a biphasic effect convergent with a hyperarousal-avoidance mechanism, as previously reported for an overlapping early time interval also at dorsal PFC regions (Wessing et al., 2017). However, this interpretation is clearly post-hoc, and more research is needed to clarify whether this region within this early time interval has excitatory or inhibitory effects on rational choice behavior. Second, we observed a significant cluster for the three-way interaction of stimulation by frame by decision in right prefrontal and temporal regions ranging from 470 to 600ms (*p*-cluster = 0.022). This cluster could be assessed as a modification of the preceding (220–270ms) significant cluster with an interaction of stimulation by frame in similar right anterior temporal/orbitofrontal areas (see Fig. 1C main text). While the pattern of the two-way stimulation by frame interaction in the ‘keep’ condition (left side of Fig. SM6) was very similar to the preceding stimulation by frame interaction (see Fig. 1C main text), the pattern was rather opposite in the ‘gamble’ condition (right side).

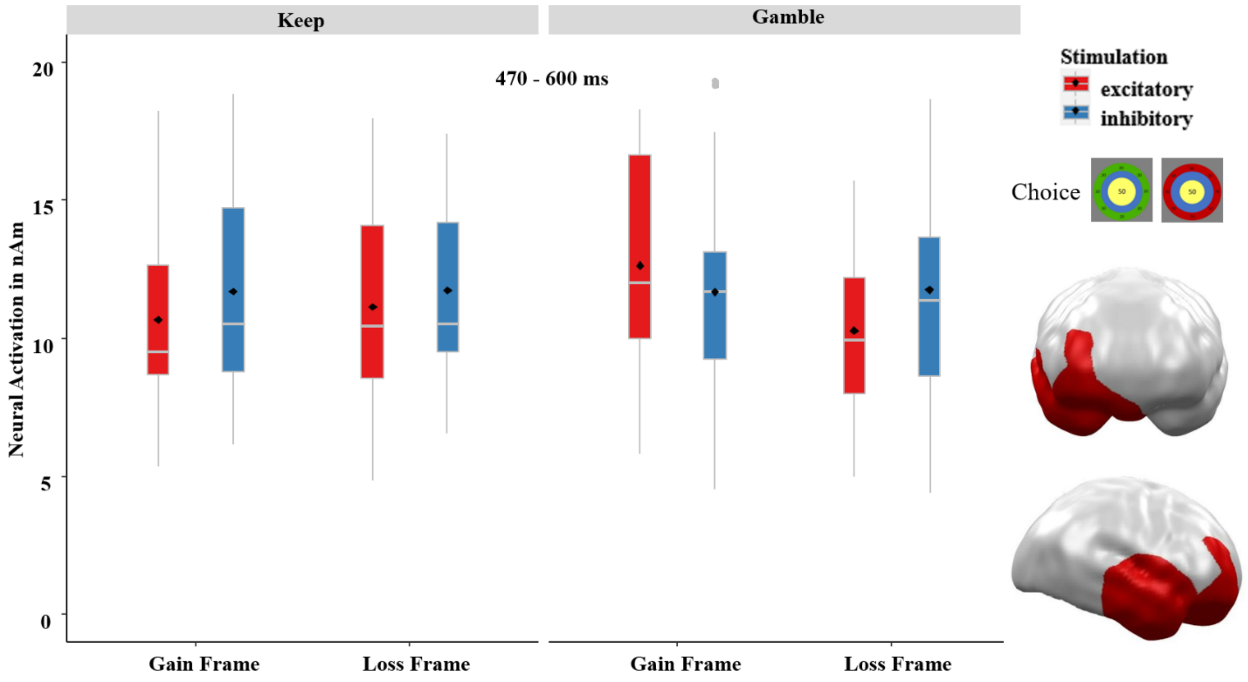


*Figure SM7.* Significant spatio-temporal cluster in right prefrontal and temporal areas featuring a three-way interaction of stimulation by decision by frame revealed by an ANOVA. Topographies of effects observed in L2-MNE were projected on standard 3D brain models for visualization Boxplots indicate means (black dots), medians (grey lines) and lower and upper quartiles. Asterisks indicate significance levels: + < 0.1, * < 0.05, ** < 0.01, *** < 0.001.

After excitatory stimulation, the difference in the loss-framed option between ‘keep’ and ‘gamble’ is smaller, which might be a sign for more elaborate processing and greater ability to inhibit irrational gambling in the loss-framed option after excitatory stimulation. The opposite pattern is present in the gain-framed option, whereby greater activity occurred if ‘gamble’ was chosen after excitatory stimulation, which possibly suggests enhanced weighing of the eventually more lucrative ‘gamble’ option, while inhibitory stimulation induces an impulsive selection of the ‘keep’ option. The pattern of the ‘keep’ condition is in line with the cluster in Fig. 1C (see main text), leading to the conclusion that the present cluster might involve temporal continuation with a more elaborated component that differentiates between ‘keep’ and ‘gamble’ decisions. Nevertheless, this three-way interaction is quite difficult to interpret, and more research is needed to replicate and illuminate this complex differential impact of stimulation on choice behavior in this late time interval.

##### 2.3.5. Correlation of neural activity and rationality

In an exploratory analysis, we correlated the overall expected value of performed decisions (i.e., the overall winnings, which could be interpreted as rationality index) of participants with their neural activity in response to the choice stimulus. The analysis revealed a cluster in left and medial prefrontal regions that emerged at 120–390ms ($p$-cluster = 0.006; $r(27) = 0.59$, $p < 0.001$). Following excitatory stimulation, the correlation was slightly stronger ($r(27) = 0.58$, $p < 0.001$) compared to the inhibitory condition ($r(27) = 0.52$, $p = 0.003$), although they did not differ significantly from each other ($z = 0.29$, $p = 0.380$).

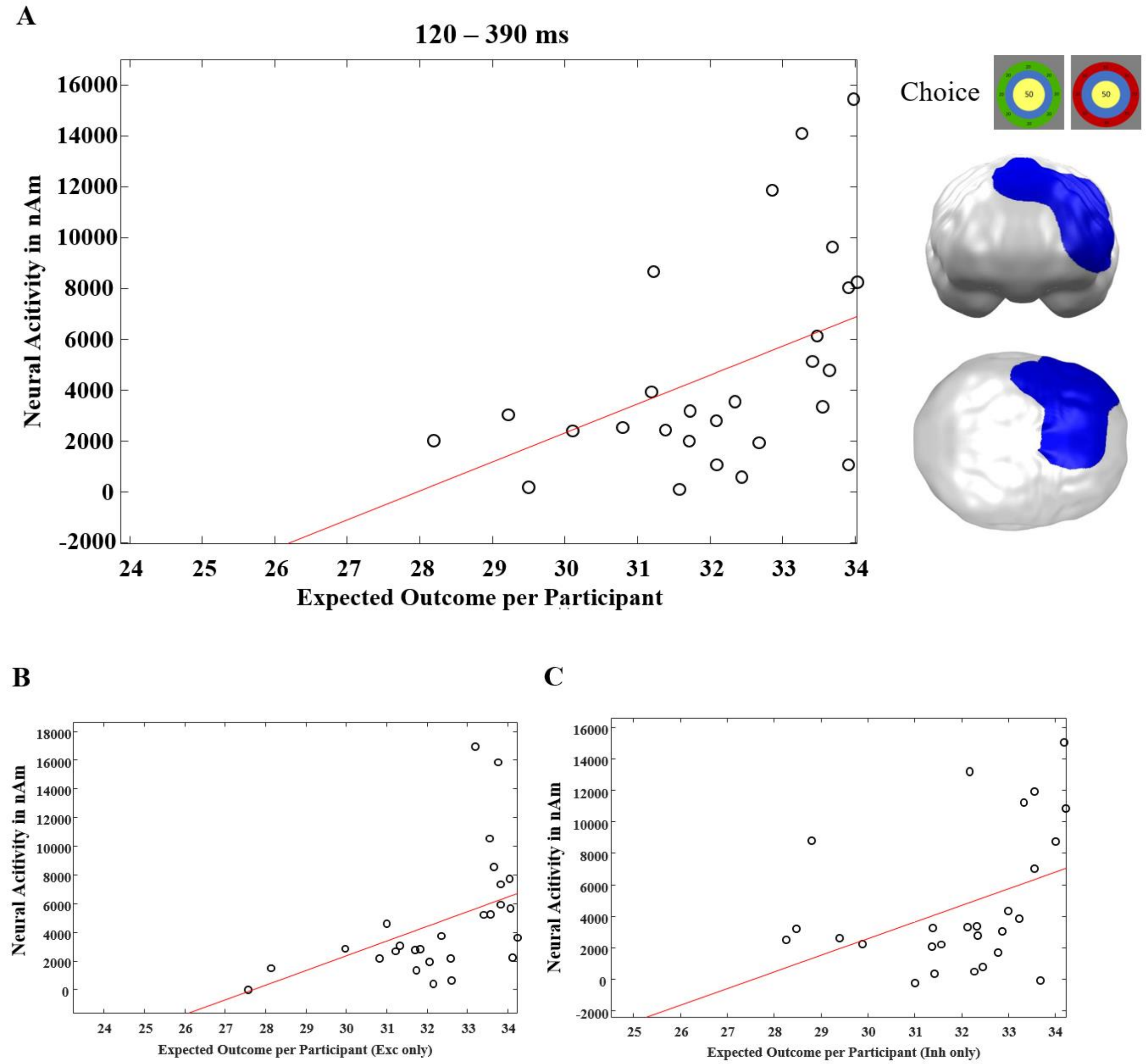


*Figure SM8.* **A.** Significant spatio-temporal cluster in left and medial prefrontal areas featuring a correlation of neural activity with the expected value of choice (i.e., overall winnings as correlate of rationality of choice behavior) The greater the neural activation in this cluster, the greater the expectation value of the decision made. **B.** Correlations in the cluster shown in A. but after excitatory stimulation only. **C.** Correlations in the cluster shown in A. but after inhibitory stimulation only. Following excitatory stimulation, the correlation was slightly stronger compared to the inhibitory condition, although they did not differ significantly from each other. Topographies of effects observed in L2-MNE were projected on standard 3D brain models for visualization.

This correlation between neural activity and the expected value of the decisions made, what can be interpreted as an index of rationality of decision-making, further supported the relevance of left and medial prefrontal regions in rational decision-making. This cluster (Fig. SM8) strongly overlaps with the subsequent (470–510ms) cluster shown in Fig. 3C (see main text), which has been shown as associated with rationality in feedback-processing, further underscoring our finding that the medial prefrontal cortex is responsible for rational evaluation of options for action and feedback on the performed decision. Again, while the later cluster in the main text (see Fig. 3C) revealed an interaction with the factor stimulation, activity in this earlier cluster shown here was not modulated by stimulation. It thus remains to be resolved if the stimulation had no sufficient impact to prove a modulation on the rationality of decision-making also in this earlier time interval.

#### 2.4. Supplementary analyses on feedback-processing

##### 2.4.1. SAM-arousal ratings

The SAM-arousal ratings did not reveal any main ($F(1,35) = 0.38$, $p = 0.538$) or interaction effects with the factor stimulation (stimulation by decision: $F(1,35) = 0.08$, $p = 0.775$; stimulation by outcome: $F(1,35) = 1.57$, $p = 0.219$; stimulation by decision by outcome: $F(1,35) = 0.12$, $p = 0.730$). Unsurprisingly, the main effect of decision was highly significant, with ‘gamble’ decisions being rated more arousing than ‘keep’ decisions $F(1,35) = 113.30$, $p < 0.001$). The main effect of outcome was insignificant ($F(1,35) = 0.51$, $p = 0.482$), while the interaction effect of decision by outcome was significant ($F(1,35) = 9.89$, $p = 0.003$). Gains were rated as more arousing after ‘gamble’ decisions, and losses were rated as more arousing in response to ‘keep’ decisions.

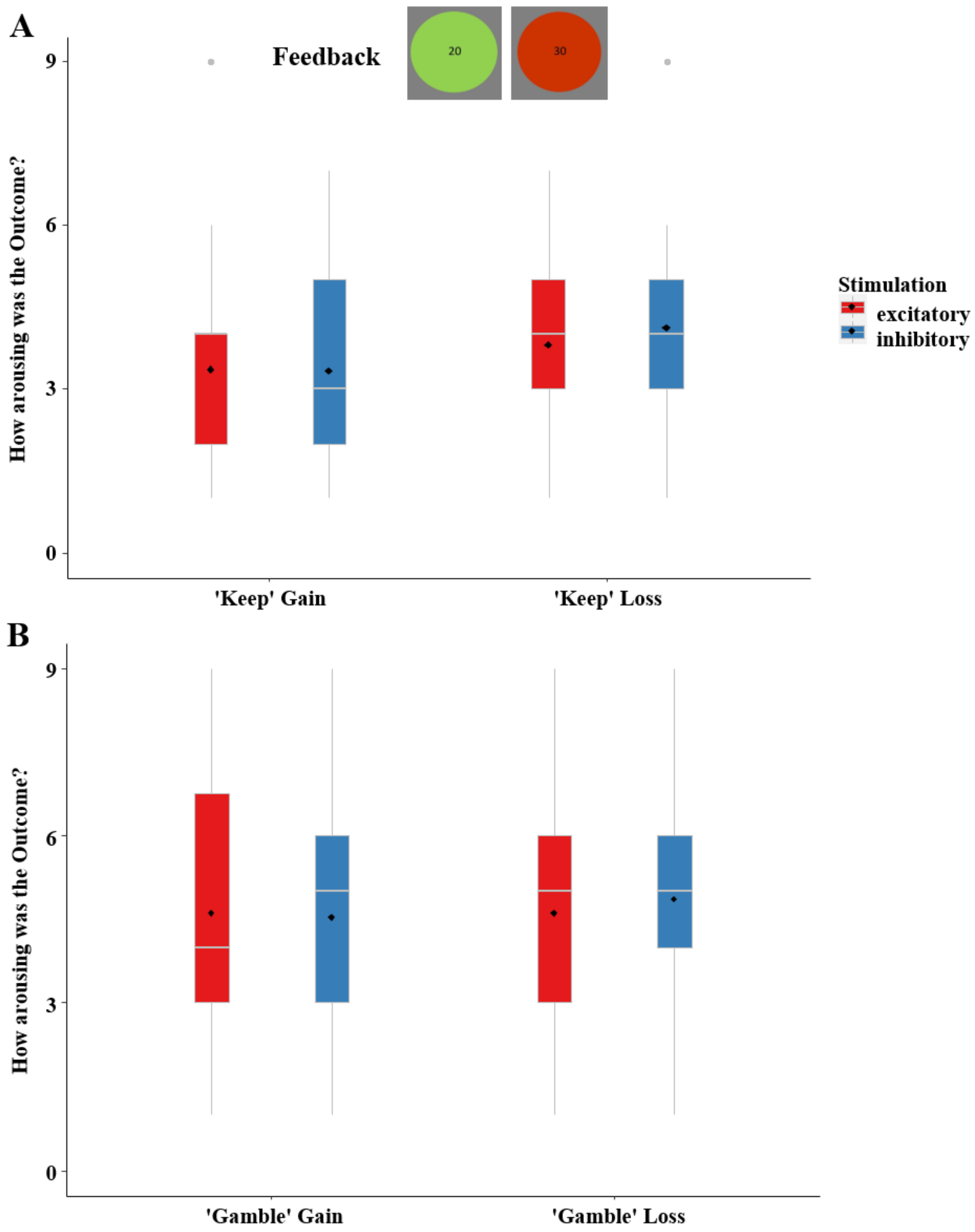


*Figure* SM9. SAM-ratings of emotional arousal (1 = not arousing at all, 9 = highly arousing) in dependency of stimulation and outcome. After ‘keep’ decisions (**A.**) losses were rated as more arousing than gains, while ‘gamble’ decisions (**B.**) evoked the opposite pattern. Boxplots indicate means (black dots), medians (grey lines) and lower and upper quartiles. Asterisks indicate significance levels: + < 0.1, * < 0.05, ** < 0.01, *** < 0.001.

##### 2.4.2. Behavioral interaction of stimulation and decisions

With our specific interest regarding the interaction of stimulation by decision, we calculated a repeated-measures ANOVA to analyze the perceived hedonic valence (SAM rating) of outcomes. This term was insignificant in the mixed-effects linear regression (see main text). Nevertheless, the ANOVA indicated a significant interaction of stimulation by decision ($F(1,35) = 20.54$, $p < 0.001$), whereby the 'keep' option was evaluated more positively than the 'gamble' option after excitatory tDCS, with an opposite pattern in the inhibitory condition.

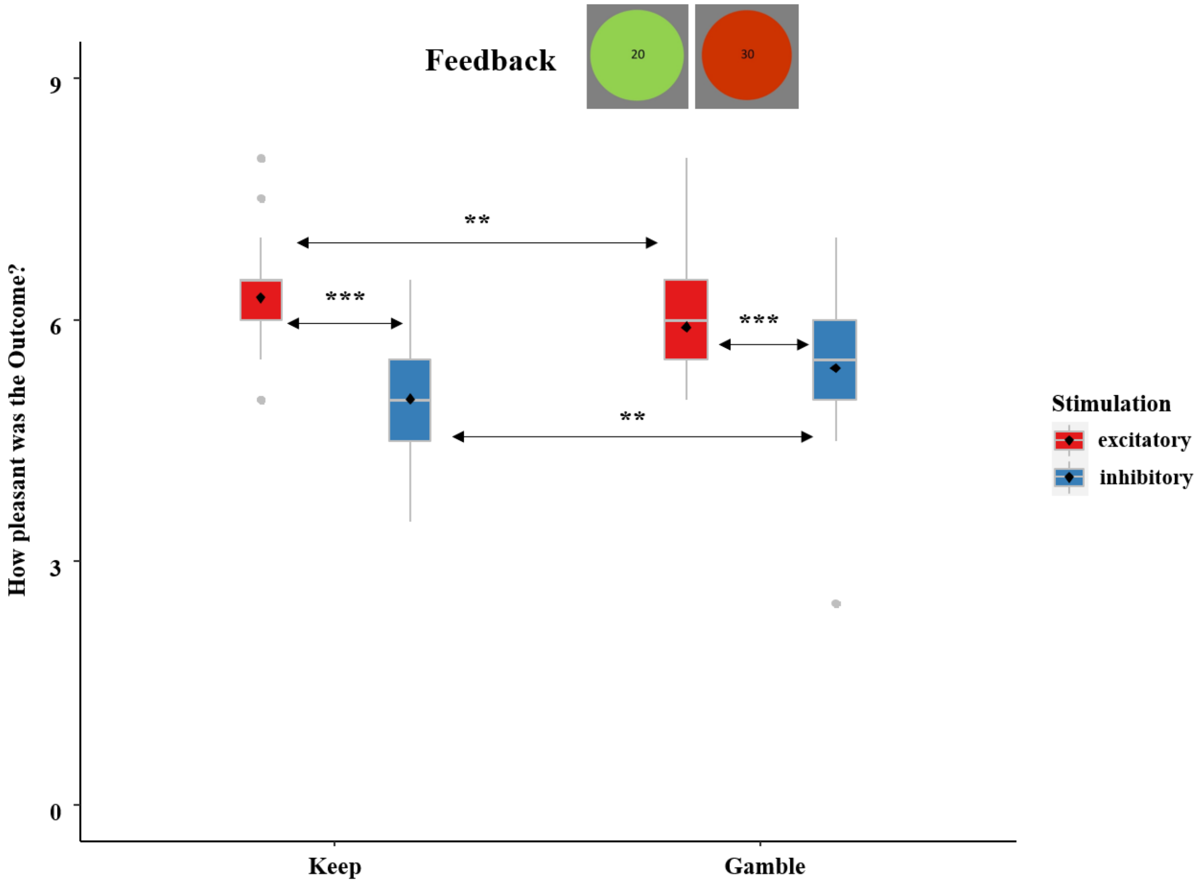


*Figure* SM10. SAM-ratings of hedonic valence (1=highly unpleasant, 9=highly pleasant) in dependency of stimulation and decision. Following excitatory stimulation, the results of the ‘keep’ option were rated more positive than the ones of the ‘gamble’ option and vice versa after inhibitory stimulation. Boxplots indicate means (black dots), medians (grey lines) and lower and upper quartiles. Asterisks indicate significance levels: + < 0.1, * < 0.05, ** < 0.01, *** < 0.001.

Since ‘keep’ outcomes were evaluated more positively than the outcomes following ‘gamble’ decisions after excitatory stimulation and, vice versa in the inhibitory condition, one can interpret this as another sign that excitatory vmPFC-tDCS allowed for a more responsible evaluation of gambling. This result matches our other behavioral results (see Fig. 2B, 2C in the main text and Fig. SM2) as well as findings of other authors (Manuel et al., 2019; Reuter et al., 2005).

##### 2.4.3. Neural main effect of outcome

Finally, we observed a main effect of outcome, that covered the complete inverse source model and got significant in the complete time interval of interest from 0[3] – 600ms ($p$-cluster < 0.001; $F(1,27) = 86.04$, $p < 0.001$). In this cluster, much stronger activations occurred in response to losses compared to gains.

[3] It should be noted that to prevent latency shifts, MEG data has been filtered in a forward-backward fashion. As backward filtering smears effects to time points preceding the real effects, statistical effects might occur even preceding the real onset of extremely strong effects.

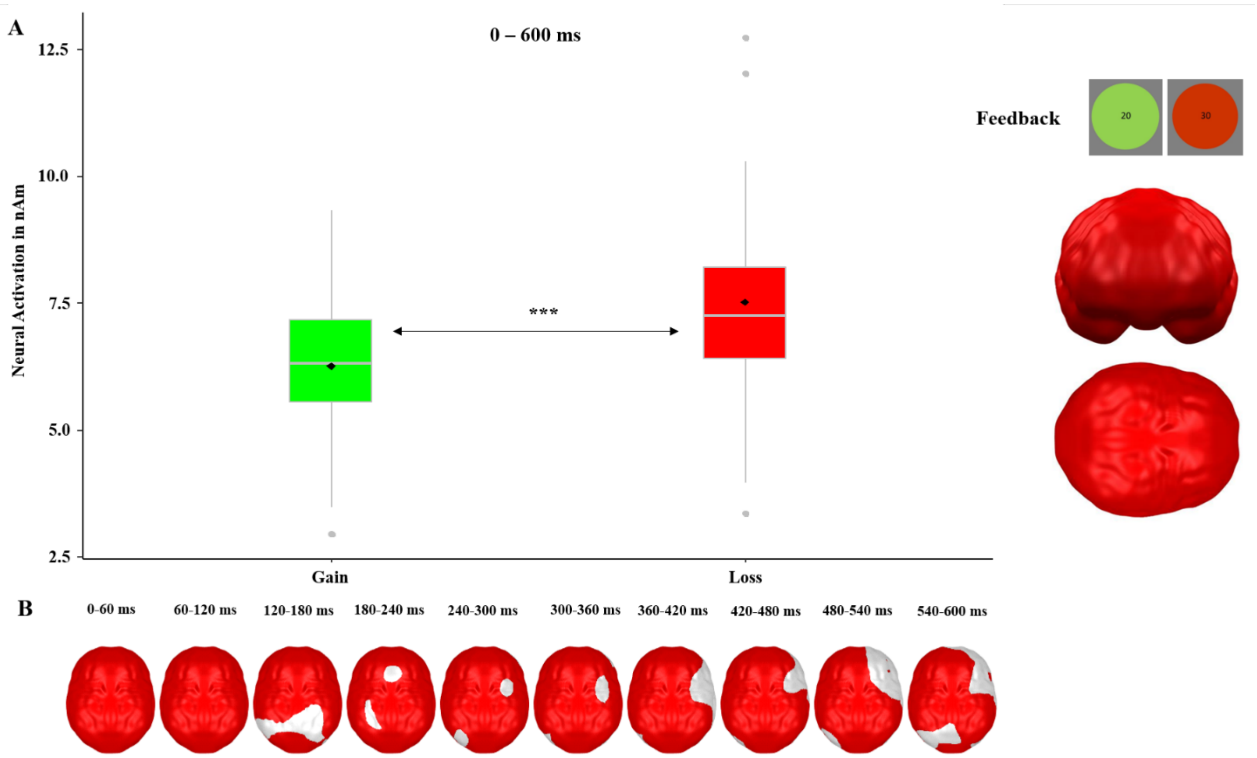


*Figure SM11.* **A.** Significant spatio-temporal cluster featuring a main effect of outcome revealed by an ANOVA. **B.** Time courses of the significant spatio-temporal cluster with a main effect of outcome revealed by an ANOVA. Boxplots indicate means (black dot), medians (grey line) and lower and upper quartiles. Asterisks indicate significance levels: + < 0.1, * < 0.05, ** < 0.01, *** < 0.001.

This cluster most likely reflects the individuals strong loss-aversion, as losses are typically rated as being around twice as salient as gains of the same value (Kahneman and Tversky, 1979; Tversky and Kahneman, 1992).

#### SM3.References

Beck, A.T., Steer, R.A., Brown, G.K., 1996. BDI-II.

Crowne, D.P., Marlowe, D., 1960. A new scale of social desirability independent of psychopathology. J. Consult. Psychol. 24, 349–354. https://doi.org/10.1037/h0047358

Gerlach, A.L., Andor, T., Patzelt, J., 2008. Die bedeutung von unsicherheits-intoleranz für die generalisierte angststörung: Modellüberlegungen und Entwicklung einer deutschen version der unsicherheitsintoleranz-skala. Z. Klin. Psychol. Psychother. 37, 190–199. https://doi.org/10.1026/1616-3443.37.3.190

Hämäläinen, M.S., Ilmoniemi, R.J., 1994. Interpreting magnetic fields of the brain: minimum norm estimates. Med. Biol. Eng. Comput. 32, 35–42. https://doi.org/10.1007/BF02512476

Junghofer, M., Winker, C., Rehbein, M.A., Sabatinelli, D., 2017. Noninvasive Stimulation of the Ventromedial Prefrontal Cortex Enhances Pleasant Scene Processing. Cereb. Cortex 27, 3449–3456. https://doi.org/10.1093/cercor/bhx073

Kahneman, D., Tversky, A., 1979. Prospect Theory: An Analysis of Decision under Risk. Econometrica 47, 263–291.

Lindquist, K.A., Satpute, A.B., Wager, T.D., Weber, J., Barrett, L.F., 2016. The Brain Basis of Positive and Negative Affect: Evidence from a Meta-Analysis of the Human Neuroimaging Literature. Cereb. Cortex 26, 1910–1922. https://doi.org/10.1093/cercor/bhv001

Manuel, A.L., Murray, N.W.G., Piguet, O., 2019. Transcranial direct current stimulation (tDCS) over vmPFC modulates interactions between reward and emotion in delay discounting. Sci. Rep. 9, 1–9. https://doi.org/10.1038/s41598-019-55157-z

Maris, E., Oostenveld, R., 2007. Nonparametric statistical testing of EEG- and MEG-data. J. Neurosci. Methods 164, 177–190. https://doi.org/https://doi.org/10.1016/j.jneumeth.2007.03.024

Reuter, J., Raedler, T., Rose, M., Hand, I., Gläscher, J., Büchel, C., 2005. Pathological gambling is linked to reduced activation of the mesolimbic reward system. Nat. Neurosci. 8, 147–148. https://doi.org/10.1038/nn1378

Roesmann, K., Kroker, T., Hein, S., Rehbein, M., Winker, C., Leehr, E.J., Klucken, T., Junghöfer, M., 2021. Transcranial direct current stimulation of the ventromedial prefrontal cortex modulates perceptual and neural patterns of fear generalization. Biol. Psychiatry Cogn. Neurosci. Neuroimaging. https://doi.org/10.1016/j.bpsc.2021.08.001

Rogers, R.D., Ramnani, N., Mackay, C., Wilson, J.L., Jezzard, P., Carter, C.S., Smith, S.M., 2004. Distinct portions of anterior cingulate cortex and medial prefrontal cortex are activated by reward processing in separable phases of decision-making cognition. Biol. Psychiatry 55, 594–602. https://doi.org/10.1016/j.biopsych.2003.11.012

Sescousse, G., Caldú, X., Segura, B., Dreher, J.C., 2013. Processing of primary and secondary rewards: A quantitative meta-analysis and review of human functional neuroimaging studies. Neurosci. Biobehav. Rev. 37, 681–696. https://doi.org/10.1016/j.neubiorev.2013.02.002

Shiv, B., Loewenstein, G., Bechara, A., Damasio, H., Damasio, A.R., 2005. Investment behavior and the negative side of emotion. Psychol. Sci. 16, 435–439. https://doi.org/10.1111/j.0956-7976.2005.01553.x

Tversky, A., Kahneman, D., 1992. Advances in prospect theory: Cumulative representation of uncertainty. J. Risk Uncertain. 5, 297–323. https://doi.org/10.1007/BF00122574

Van den Berg, I., Franken, I.H.A., Muris, P., 2010. A new scale for measuring reward responsiveness. Front. Psychol. 1, 1–7. https://doi.org/10.3389/fpsyg.2010.00239

Watson, D., Clark, L.A., Tellegen, A., 1988. Development and validation of brief measures of positive and negative affect: the PANAS scales. J. Pers. Soc. Psychol. 54, 1063–1070. https://doi.org/10.1037//0022-3514.54.6.1063

Wessing, I., Romer, G., Junghöfer, M., 2017. Hypervigilance-avoidance in children with anxiety disorders: magnetoencephalographic evidence. J. Child Psychol. Psychiatry Allied Discip. 58, 103–112. https://doi.org/10.1111/jcpp.12617

Winker, C., Rehbein, M.A., Sabatinelli, D., Dohn, M., Maitzen, J., Roesmann, K., Wolters, C.H., Arolt, V., Junghoefer, M., 2019. Noninvasive stimulation of the ventromedial prefrontal cortex indicates valence ambiguity in sad compared to happy and fearful face processing. Front. Behav. Neurosci. 13, 1–16. https://doi.org/10.3389/fnbeh.2019.00083

Winker, C., Rehbein, M.A., Sabatinelli, D., Dohn, M., Maitzen, J., Wolters, C.H., Arolt, V., Junghofer, M., Junghöfer, M., 2018. Noninvasive stimulation of the ventromedial prefrontal cortex modulates emotional face processing. Neuroimage 175, 388–401. https://doi.org/10.1016/j.neuroimage.2018.03.067

Winker, C., Rehbein, M.A., Sabatinelli, D., Junghofer, M., 2020. Repeated noninvasive stimulation of the ventromedial prefrontal cortex reveals cumulative amplification of pleasant compared to unpleasant scene processing: A single subject pilot study. PLoS One 15, e0222057.

Xue, G., Lu, Z., Levin, I.P., Bechara, A., 2011. An fMRI study of risk-taking following wins and losses: Implications for the gambler's fallacy. Hum. Brain Mapp. 32, 271–281. https://doi.org/10.1002/hbm.21015
